# Using domain insertion to create sulfite reductases that present chemical-dependent activities

**DOI:** 10.1101/2025.05.23.655854

**Authors:** Elizabeth Windham, Dru Myerscough, Samuel K. Schwartz, Matthew D Carpenter, Caroline M. Ajo-Franklin, Jonathan J. Silberg

**Affiliations:** Department of Biosciences, Rice University, 6100 Main Street, Houston, TX, 77005; Systems, Synthetic, and Physical Biology Graduate Program, Rice University, 6100 Main Street, Houston, Texas 77005; Department of Bioengineering, Rice University, 6100 Main Street, Houston, TX, 77005; Department of Chemical and Biomolecular Engineering, Rice University, 6100 Main Street, Houston, TX, 77005

**Keywords:** bioelectronics, domain insertion, extracellular electron transfer, protein engineering, oxidoreductase, sensor, synthetic biology, and sulfite reductase

## Abstract

Domain insertion can be used to create oxidoreductase switches whose charge transfer is dependent upon analyte binding. To date, most domain insertion studies have targeted relatively small proteins of known structure, so it remains unclear how to effectively use this protein engineering approach with large oligomeric oxidoreductases that require dynamic conformational changes for catalysis. To address this question, we studied the effect of domain insertion on the function of NADPH-dependent sulfite reductase (SiR) from *Escherichia coli*, a dodecameric protein containing eight hemoprotein subunits and four flavoprotein subunits. SiR mutational tolerance was first mapped using systematic peptide insertion, and a subset of variants retaining activity were subjected to domain insertion. When a ligand-binding domain was inserted at locations tolerant to peptide insertion, including sites proximal and distal from the intersubunit interfaces, more than half of the resulting variants presented cellular activity that is enhanced by an endocrine disruptor. This ligand-dependent synthesis of a redox-active metabolite could be monitored electrochemically from cells, illustrating how a single protein complex can be used to convert chemical information in the environment into a metabolite within cells that diffuses across the cell membrane and can be detected electrochemically.

## INTRODUCTION

Chemical contaminants that enter the environment pose a threat to our health and to our resources^1^. Many of these contaminants enter our environment from diverse waste streams^2^, driving a need for distributed sensors that report on when and where chemical hazards are present. Synthetic biology can be leveraged to overcome this challenge by creating biosensors^3^, living systems that convert chemical information in the environment into easy to detect signals. Biosensors are commonly programmed to report by generating visual outputs, such as the fluorescent and luminescent proteins^4,5^. To enable applications in hard-to-image settings, alternative outputs can be used, such as ice nucleation proteins^6^, indicator gases^7^, and bioelectronic reporters^8^. Among these reporters, only bioelectronic reporters generate electrical information directly without requiring additional analytical instrumentation.

Living sensors can be programmed to convert chemical information into electrical information by controlling a process called extracellular electron transfer (EET)^9^. EET has been made conditional by programming cells to regulate the expression of proteins that participate in EET^10^, including proteins that directly transfer electrons from cells to extracellular materials and proteins that control the reduction of chemical mediators that diffuse in and out of cells^11–16^. Recently, bioelectronic sensors were created by regulating EET post-translationally^17^, similar to that achieved *in vitro* with glucose dehydrogenase sensors^18–20^. This EET regulation, which was achieved in *Escherichia coli*, allowed for real-time sensing of endocrine disruptors in riverine samples^17^. To achieve this sensing, cells were programmed to express a complex, eight-component synthetic electron transfer pathway that contains a protein switch along with a dozen other proteins.

To explore if a simpler pathway design strategy can be used to create living electronic sensors, we investigated whether sulfite reductase (SiR) can be engineered to exhibit conditional activity. This oxidoreductase was targeted because it synthesizes hydrogen sulfide, a redox active metabolite that can diffuse across the cell membrane^21^. Here, we target the SiR hemoprotein (SiR-HP) for switch design, which catalyzes the six electron reduction of sulfite to sulfide^22^, and examine the activity of SiR-HP variants when coexpressed with the native SiR flavoprotein (SiR-FP), which provides reducing equivalents for catalysis through formation of a large dodecameric complex containing eight SiR-FP and four SiR-HP subunits^23^. We identify multiple SiR-HP whose activities are switched on by endocrine disruptors and show that one of these can be used in *E. coli* to create a bioelectronic sensor.

## RESULTS

To assay sulfide production by SiR, we evaluated growth complementation of an *E. coli* strain (EW11) that lacks a functional SiR^24^. This strain was able to grow in medium having sulfate as the only sulfur source when it transcribed the *cysJI* operon using the TetR promoter (Supplementary Fig. 1). However, growth was not observed when cells expressed the SiR hemoprotein (SiR-HP) having an inactivating R83A mutation. The *cysJI* vector that complemented cell growth was used to create a library of SiR-HP mutants having the octapeptide SGRPGSLS inserted at each backbone location (Supplementary Figs. 2a-b). This peptide was chosen because it arises from the insertion of a unique restriction site that allows for facile domain insertion^25^. Next generation sequencing (NGS) revealed that 560 out of 570 possible insertion variants (98%) were present in the library (Supplementary Fig. 2c). A mixture of *E. coli* EW11 harboring this library and vectors expressing SiR-HP with a silent barcode or with the inactivating R83A mutation was used to inoculate selective and unselective cultures (Supplementary Fig. 3a). In selective conditions, the cultures seeded with the library grew to a higher density compared to those seeded with cells expressing the R83A mutant (Supplementary Fig. 3b). This latter finding suggests that a subset of the SiR-HP peptide insertion variants in the library are active.

NGS analysis revealed similar sequence diversity across the three biological replicates performed with the naïve (Supplementary Fig. 4), unselected (Supplementary Fig. 5), and selected (Supplementary Fig. 6) libraries. A comparison of average sequence abundances revealed that the selected library presents a greater dispersion of average sequence abundances than the naïve and unselected libraries (Supplementary Fig. 7).

To identify which SiR-HP variants retain activity, the sequence enrichment of each variant was compared with the embedded controls (Fig. 1a). This enrichment data could be described by a gaussian mixture model having two peaks whose values are similar to the controls (Fig. 1b). When the enrichment scores were mapped onto the monomeric SiR- HP monomer structure (Fig. 1c), peptide-insertion tolerance was found to be significantly higher in loops compared with other secondary structure (Supplementary Fig. 8a). In addition, insertion tolerance was generally inversely correlated with distance from the siroheme and 4Fe-4S cofactors (Supplementary Fig. 8b). Surprisingly, two loops proximal to the active site (≤10Å) presented enrichment scores implicating those variants as functional (Supplementary Fig. 8c). These trends are similar to other peptide insertion studies, which have found that loops present high insertion tolerance and cofactor proximity is inversely correlated with insertion tolerance^26–28^.

**Figure 1.**
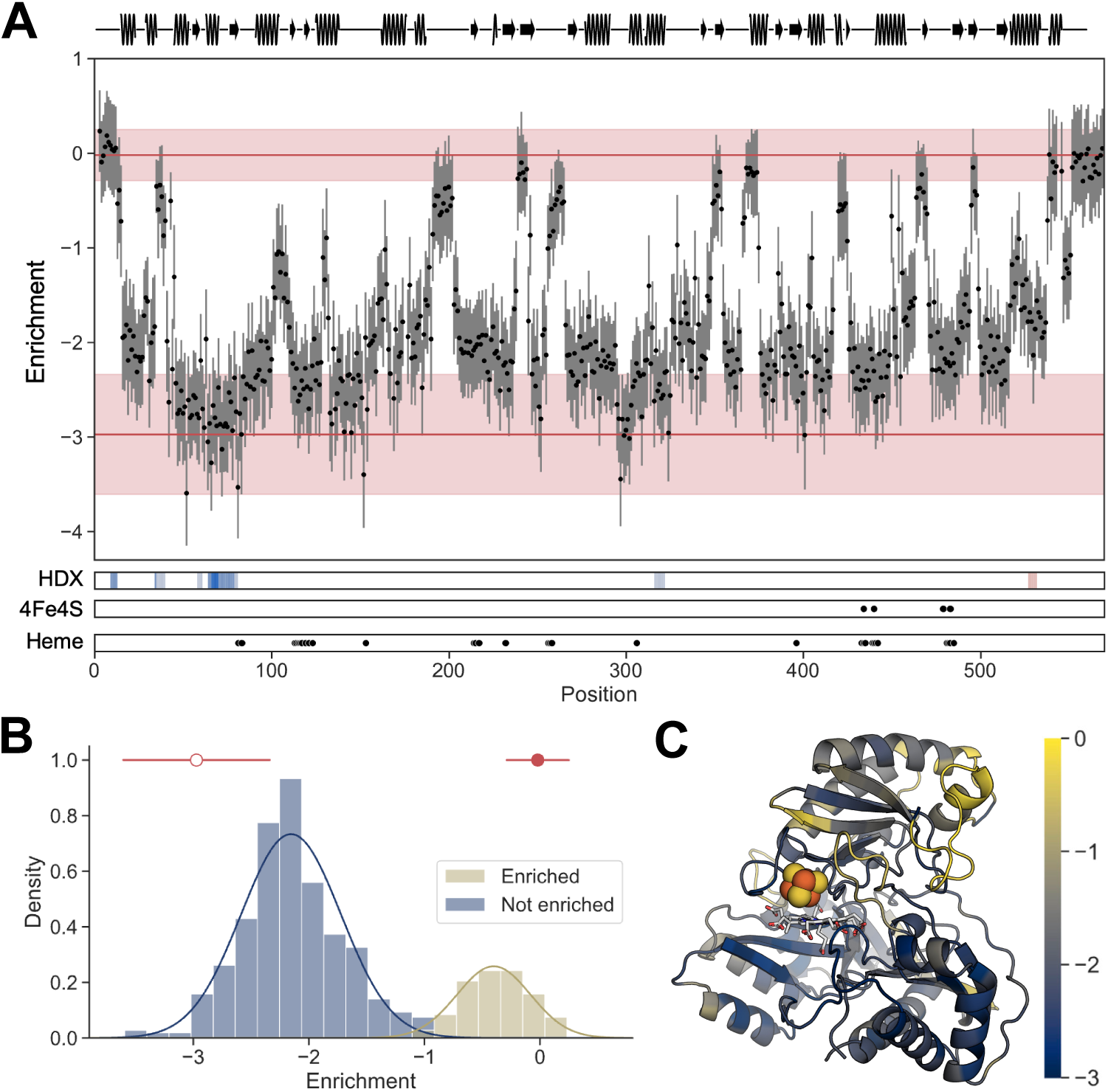
SiR-HP tolerance to peptide insertion. (**A**) The enrichment value for each peptide-insertion variant is mapped onto primary structure. These values were calculated by taking a ratio of the selected library reads from three biological replicates to the naïve library reads and using the Rosace program to calculate the enrichment values. Each dot point represents the enrichment value for a single variant with the gray lines being two standard deviations. The upper red line is the enrichment value for the embedded positive control with the shaded region being ±1 standard deviation, and the lower red line is the same for the negative control. The data is compared with SiR-HP secondary structure (top), residues that coordinate the 4Fe-4S cluster (4Fe4S) and contact the siroheme (heme), and prior H-D exchange data (HDX); this latter data shows regions that are more buried (blue) in the SiR-HP/FP multimer compared with the SiR-HP monomer and residues that are less buried (red). (**B**) The bimodal distribution of the enrichment values fit to a two-component guassian mixture model. The open and closed circles represent the average enrichment values for the negative and positive controls, respectively, with lines representing ±1 standard deviation. (**C**) The monomeric structure of SiR-HP (PDB 6c3m)^32^ is colored according to enrichment. Yellow represents high enrichment, while blue represents low enrichment.

Mutant enrichment was also mapped onto the SiR-HP/FP heterodimer structure (Supplementary Fig. 9a), which was recently obtained using cryoelectron microscopy^29^, as well as the SiR-HP residues implicated in binding the Fld domain in SiR-FP (Supplementary Fig. 9b). To identify the latter residues, AlphaFold2 was used to generate a model for the Fld domain binding to SiR-HP^30^. In the predicted structure, the intercofactor distance between the FMN in the Fld domain of SiR-FP and the 4Fe-4S cluster in SiR-HP was 8.8 Å (Supplementary Fig. 10 *top*), which is shorter than the 12 to 14 Å distance between cofactors in ferredoxin-dependent SiRs^31^, and methionine 444 in SiR-HP, a residue implicated in gating access for electron transfer^32^, is proximal to both cofactors (Supplementary Fig. 10 *bottom*). Approximately half a dozen peptide insertions proximal to SiR-HP residues making intermolecular contacts with the SiR-FP FNR domain present high enrichment values (Supplementary Fig. 9c). When the enrichment scores at this interface in the dimer were compared with prior results from HD exchange studies utilizing the full SiR dodecamer (Supplementary Fig. 11), only a subset of the residues that presented low HD exchange at this interface present low enrichment values upon peptide insertion. Among the SiR-HP residues implicated in binding to the dynamic Fld domain in SiR-FP, which mediates ET, more than a dozen present high enrichment values (Supplementary Fig. 9d). These findings suggest that some peptide insertions are tolerated proximal to both SiR-HP/FP interfaces and do not disrupt the catalytic activity of this protein complex.

We posited that backbone locations tolerant to peptide insertion may represent locations where larger domains can be inserted to create ligand-dependent switches. To test this idea, we inserted the estrogen receptor ligand binding domain (LBD) at sites both proximal and distal to the SiR-HP and SiR-FP interfaces (Fig. 2a, Supplementary Table 1). Biophysical studies have shown that the termini of the estrogen receptor LBD are distal in the absence of ligands, but then are brought together upon ligand binding^33,34^. When assayed for sulfide production in *E. coli* EW11, a range of phenotypes were observed with these domain insertion variants (DVs) (Supplementary Fig. 12). More than half of the variants presented higher growth rates (Fig. 2b) and cell densities (Supplementary Fig. 13) in the presence of the endocrine disruptor 4-hydroxy-tamoxifen (4-HT). The most dramatic switching phenotypes were observed when the LBD was inserted after SiR residues 38 (DV38), 39 (DV39), 259 (DV259), and 543 (DV543). In unselective growth medium, growth was similar ±4-HT (Supplementary Fig. 14). These results show many sites that tolerate peptide insertion also tolerate domain insertion, and they suggest that insertion proximal to protein-protein interfaces represents an effective strategy for creating SiR-HP switches.

**Figure 2.**
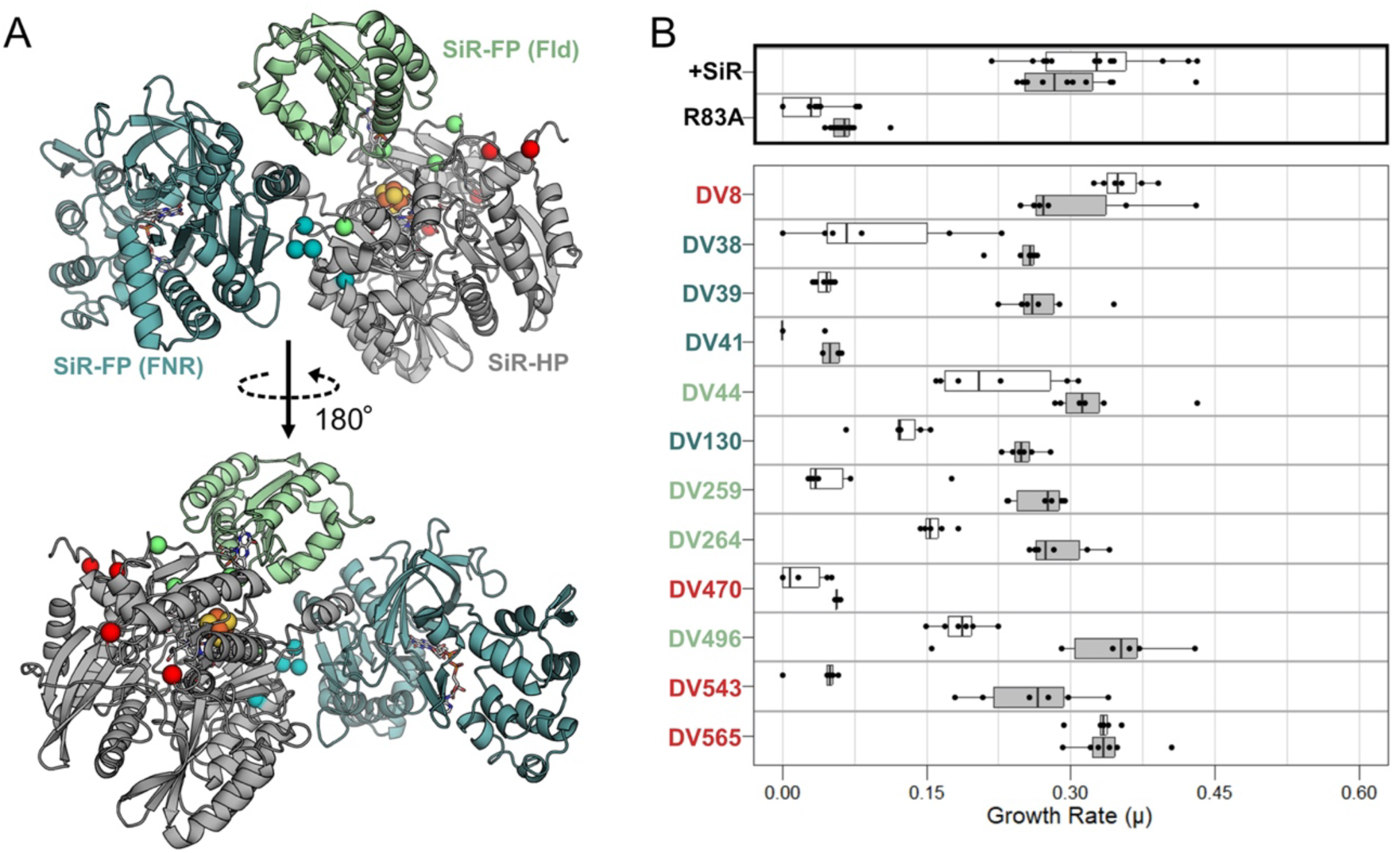
Effect of substituting a domain insertion for a peptide insertion. (**A**) The ER LBD was inserted at SiR-HP backbone locations (spheres) proximal to the interface with the SiR-FP Fld domain (green), the SiR-FP FMN domain (cyan), and distal from both interfaces (red). (**B**) (*top*) Effect of 10 µM 4-HT (gray) and the DMSO vehicle (white) on the growth complementation of *E. coli* EW11 expressing SiR-HP and SiR-FP (+SiR) or a R83A mutant of SiR-HP and SiR-FP (R83A). (*bottom*) Effect of 4-HT and DMSO on the growth complementation of *E. coli* EW11 expressing SiR-FP and SiR-HP having the ER-LBD inserted in locations that yielded high enrichment scores (>1.5). For all experiments, six biological replicates were performed (dots). Within each box, the center line represents the 50^th^ percentile, while ends of the box represent the 25^th^ and 75^th^ percentiles, respectively. Outside the box, the line tips represent the minima and maxima. Dots outside of the lines correspond to outliers. The growth of SiR-HP and domain variants 8, 41, 470, and 565 were not enhanced by 4-HT (p > 0.05, a one-tailed, two sample paired t-test), whereas domain variants 39, 44, 130, 259, 264, 496, and 543 had significantly enhanced growth (DV39, p = 7.75e-06; DV44, p = 0.026; DV130, p = 0.00039; DV259, p = 0.00059; DV264, p = 0.00023; DV496, p = 0.010; DV543, p = 0.00016, using a one-tailed, two sample paired t-test).

Estrogen receptor agonists and antagonists elicit distinct conformations and cellular responses^33,34^, suggesting that they may have different effects on SiR switch activation. To investigate this idea, we measured the growth complementation of *E. coli* EW11 expressing DV39 or DV259 (Fig. 3a) in medium containing either an agonist (17β- estradiol, diethylstilbestrol, or hexestrol), antagonists (raloxifene, lasoxifene), endocrine disruptor (bisphenol-A), or ligand vehicle (DMSO). These variants were chosen because they arise from domain insertion proximal to the structural and dynamic SiR-HP/SiR-FP interfaces, respectively. The endocrine antagonists activated sulfide production like 4-HT, while the agonists and bisphenol-A could not be differentiated from the DMSO control (Fig. 3b and Supplementary Fig. 15). These chemicals had no effect on the complementation of *E. coli* EW11 by native SiR or the R83A mutant. This finding shows that these two switches are specific for estrogen receptor antagonists, which have been shown to promote a distinct conformation from agonists^33,34^.

**Figure 3.**
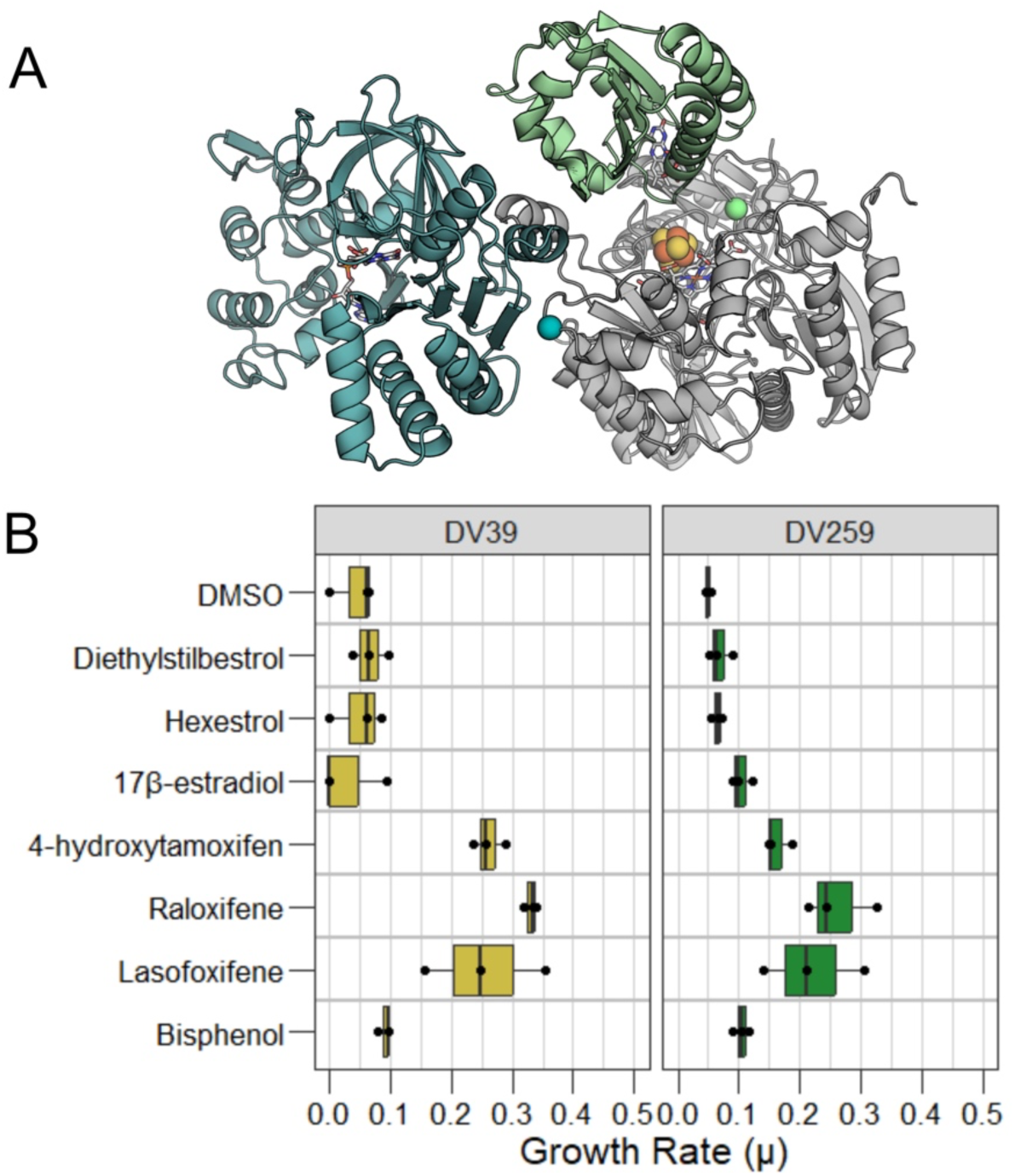
Estrogen antagonists specifically activate SIR growth complementation. (**A**) The ER-LBD insertion sites used to create DV39 (teal) and DV359 (green) are mapped as spheres onto the SiR-FP/HP heterodimer. (**B**) Growth complementation was measured in the presence of eight different chemical treatments using cells that coexpress SiR-FP and either the native SiR- HP (+SiR), the inactive R83A SiR-HP, SiR-HP domain insertion variant 39 (DV39), and SiR-HP domain insertion variant 259 (DV259). For all experiments, three biological replicates were performed (dots). Within each box, the center line represents the 50^th^ percentile, while ends of the box represent the 25^th^ and 75^th^ percentiles, respectively. Outside the box, the line tips represent the minima and maxima. Dots outside of the lines correspond to outliers. With DV39, lasofoxifene, raloxifene, and 4-hydroxytamoxifen significantly increasing growth rates (lasofoxifene, p = 0.003; raloxifene, p = 5e-6; 4-hydroxytamoxifen, p = 0.0002; post-hoc Dunnett’s test) compared to DMSO. In contrast, bisphenol, 17β-estradiol, hexestrol, and diethylstilbestrol had no effect on growth (p > 0.05, post-hoc Dunnett’s test) compared to the DMSO control. With DV259, bisphenol, 17β-estradiol, hexestrol, and diethylstilbestrol did not affect growth (p > 0.05, post-hoc Dunnett’s test) compared to DMSO, whereas lasofoxifene, raloxifene, and 4- hydroxytamoxifen increased growth (lasofoxifene, p = 0.0003; raloxifene, p = 2e-6; 4- hydroxytamoxifen, p = 0.01, post-hoc Dunnett’s test).

To investigate if a SiR switch can be used to directly couple detection of a chemical in the environment into an electrical signal, we analyzed the current produced by EW11 cells expressing DV39 using a bioelectrochemical system (Fig. 4a). With cells expressing the switch, 4-HT exposure yielded significantly higher current density compared with cells lacking 4-HT over the full time-course of the measurements (Fig. 4b). In addition, the current in the absence of 4-HT was similar to that of the R83A mutant over the time course of the experiment. However, the activity of the switch +4-HT was significantly lower than the current observed with an unmodified SiR-HP. Also, the current from this positive control presented greater variability than the switch measurements (Supplementary Fig. 16). This variability appeared to arise from variation in cell growth in one positive control replicate (Supplementary Fig. 17). These findings show that a SiR-HP switch can be used in *E. coli* to report on chemical sensing by producing an electrical signal.

**Figure 4.**
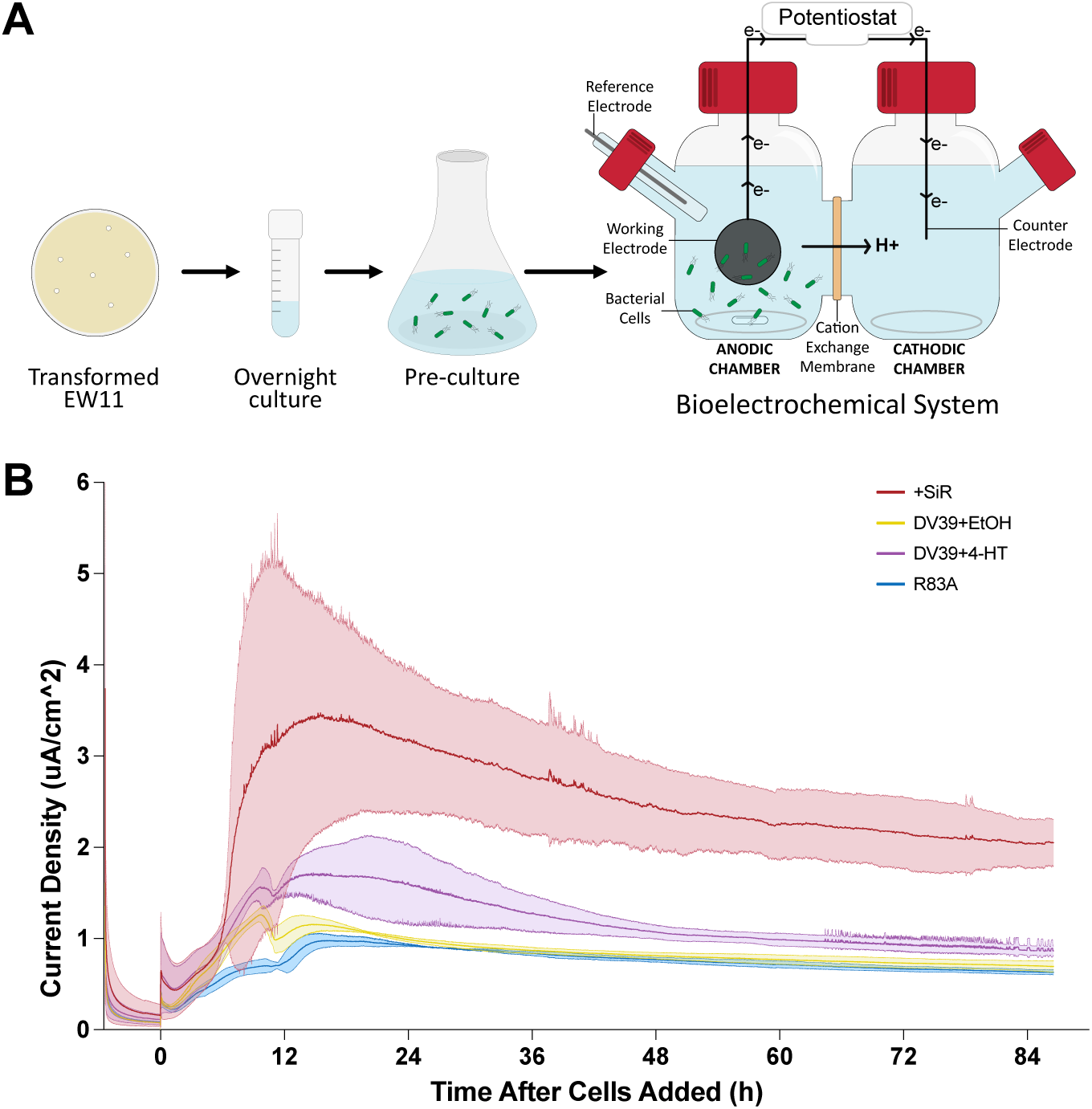
Electrochemical detection of switch-dependent sulfide production. (**A**) Workflow for electrochemical measurements in a two-chamber bioelectrochemical system (BES) reactor. *E. coli* EW11 colonies were grown through a two-stage process to obtain culture that was used to inoculate the BES at an OD of 0.2. In this BES, the *E. coli* were in the anodic chamber having a carbon felt working electrode. A potentiostat was used to measure electron flow from the working electrode through the circuitry and into the cathodic chamber^67^. (**B**) The average current density from three biological replicates is shown (lines) with standard error (shaded regions). Experiments were performed using cells expressing SiR-FP and SiR-HP with ethanol (black), DV39 with 4-HT in ethanol (blue), DV39 with ethanol (pink), and the SiR R83A mutant with ethanol (orange). Time zero represents the injection of cells, thiosulfate, and either ethanol or 4-HT in ethanol. The signal from DV39 with 4-HT is significantly higher than DV39 with ethanol only (one-tailed t-test, p = 0.03) and the negative control with ethanol (p = 0.02) at 24 hours. However, the DV39 switch treated with 4-HT is significantly lower than the positive control with ethanol only (p = 0.02).

## DISCUSSION

Our results add SiR to the list of oxidoreductases whose activities can be made chemical dependent, which includes glucose oxidase^18–20^ and ferredoxin^35–37^. Like ferredoxin switches^38^, SiR switches can be used in cells to report on analytes electrochemically. However, to generate a cellular signal, the SiR system only requires two proteins be expressed, SiR-FP and an SiR-HP switch. In contrast, pathways leveraging ferredoxin switches require designs containing twenty proteins^38^. Although the SiR system is simpler, the temporal response is slower, requiring hours for a signal. The reason for the slow kinetics is not known. It could arise because the SiR switches require ligand binding to fold and accumulate in cells^39^, such that SiR activity is limited by translation, or because the H2S that they synthesize is consumed by other reactions. To improve, future studies should explore alternative SiR switches for regulating H2S production. Also, they should investigate if the signal can be improved by tuning switch expression, deleting H2S consuming enzymes^40^, expressing transporters for sulfur anions^41^, and using electrodes that are designed for more sensitive H2S detection^42^.

With SiR, peptide insertion tolerance was a useful guide for identifying domain insertion sites that present switching activities. Among the SiR peptide insertion variants subjected to domain insertion, ∼90% retained SiR activity following ER LBD insertion and >50% presented ligand-dependent activity. This approach can be contrasted with prior systematic studies of domain insertion in GFP^43^ and CRISPR-Cas9^44^, which yielded a smaller fraction of tested designs that retained activity. Ion channels have also been systematically modified using both peptide and domain insertion at every backbone location^45^. With ion channels, peptide and domain insertion tolerance were similar at many insertion locations, although some backbone sites tolerated domain but not peptide insertion. These findings highlight an opportunity to create models for domain insertion that train on natural and synthetic insertion events of varying sizes^46^. Such models could be used to further diversify the number of electrochemical channels that report on analytes post-translationally by targeting enzymes that synthesize alternative mediators, such as enzymes that synthesize L-DOPA^47^, flavins^48^, quinones^49^, and pyocyanin^50^.

A comparison of insertion tolerance with SiR structure reveals two surprising results. First, although residues involved in cofactor binding and proton transfer present low insertion tolerance^22,32,51^, ligand-inducible activity was observed with a domain insertion variant (DV259) bearing an insertion <6 Å from the siroheme cofactor. This observation is reminiscent of prior observations that some active site mutations can increase SiR activity^52^. Further biochemical studies will be required to determine if insertion variants with apparently high activity display more subtle defects, such as the release of partially reduced products^51^. Second, a high tolerance to peptide insertion was observed at the SiR-HP and Fld domain binding interface, which mediates electron transfer from SiR-FP to SiR-HP. This observation could reflect compensatory electron transfer from adjacent SiR-FP copies in the dodecamer^23^ or that conformational changes or another process, rather than binding, may be rate-limiting for activity *in vivo*. Presumably, the peptide or domain insertions herein may disrupt activity by interfering with folding, oligomerization, electron transfer, or catalysis. Our finding of ligand-dependent activity in SiR-HP variants bearing insertions in regions associated with each of these behaviors suggests the exciting possibility of enabling artificial control of oligomerization, electron transfer, and catalysis within a multimeric assembly.

## METHODS

### Materials

Chemicals were from Fisher Scientific, Millipore Sigma, Sigma-Aldrich, ThermoFisher, and VWR. Enzymes were from New England Biolabs. Kits for molecular biology were from Qaigen and Zymo Research.

### Strains

*E. coli* XL1-Blue Electroporation-Competent Cells (Agilent) were used for vector construction, *E. coli* MegaX DH10B T1 Electrocomp Cells (ThermoFischer Scientific) were used for library construction, and *E. coli* EW11 were used for growth complementation^24^. Cells were cultured in lysogeny broth (LB), unless otherwise noted. **Plasmids.** Table S2 lists all vectors used. To create a vector that complements the growth of *E. coli* EW11, the *cysJI* operon from *E. coli*, including the native translation initiation sites, was PCR amplified from *E. coli* MG1655 genomic DNA and cloned into plasmid containing the TetR promoter using Golden Gate Assembly^53^. The resulting plasmid (pLW001) has a ColE1 origin and chloramphenicol resistance (Cm^R^). To create a positive control vector (pLW002) for library selections, a 14 base pair barcode was inserted after the *cysJI* terminator in pLW001. To create a negative control vector (pLW003), a CysI mutation (R83A) was incorporated into pLW001, which inactivates SiR-HP^51^. To create vectors that express SiR containing an inserted ER-LBD domain, a gBlock was synthesized that encodes ER-LBD residues 6 to 256 flanked by sequences coding for residues (GGGGSGGGGS) that would ultimately code for linkers in domain insertion variants (Integrated DNA Technologies). This gBlock was cloned into vectors having peptide insertions, which code for a unique BsaI cloning site, using Golden Gate Assembly^53^. All vectors were sequence verified (Plasmidsaurus Inc).

### Library construction

The peptide insertion library was designed using the SPINE algorithm^25^, which divided the SiR-HP gene (*cysI*) into 10 tiles having 55 to 60 codons, and predicted an oligonucleotide pool for library construction having the sequence AGCGGGAGACCGGGGTCTCTGAGC inserted after every codon. Commercially synthesized tiles (Twist Biosciences) were PCR amplified using primers designed by SPINE^25^ and cloned into pLW001 using Golden Gate cloning^53^. The product of these reactions was concentrated using DNA Clean & Concentrator-5 kit (Zymo Research) and transformed into *E. coli* MegaX Electrocomp Cells. After recovery at 37°C for 1 hour while shaking at 250 rpm, cells were plated on LB-agar medium containing chloramphenicol (34 µg/mL). Following incubation at 37°C overnight, plates from each cloning reaction (n = 55 to 60 variants) contained ≥1000 CFU. This CFU count was estimated to cover >99.9% of the variants possible in each tile assuming sampling with replacement^54^. Colonies were harvested, and plasmid was purified from using a Qaigen Miniprep kit. To assess library coverage, each tile was sequenced using NGS (Azenta, Inc) To create the naïve library, plasmids arising from each cloning reaction were mixed at stoichiometric ratios.

### Growth complementation

To assay SiR activity, the growth of *E. coli* EW11 was evaluated on selective growth medium containing sulfate as the only sulfur source (M9sa). As an unselective growth medium (M9c), M9sa was modified to contain cysteine and methionine as a sulfur source (M9c). M9c and M9sa contained M9 salt stock, 2% glucose, tryptophan (40 mg/L), tyrosine (1.2 mg/L), all other amino acids lacking sulfur (80 mg/L each), vitamins stock, calcium chloride (11 mg/L), ferric citrate (24 mg/L), and magnesium sulfate (1.2 g/L). M9sa additionally contained cysteine (80 mg/L) and methionine (80 mg/L). The vitamin stock contained p-aminobenzoic acid, inositol, adenine and uracil. To assess growth complementation, single colonies obtained from LB-agar were grown in LB medium (3 mL) containing chloramphenicol. After overnight growth, cultures were washed two times and resuspended in M9sa medium (1 mL). The optical density (OD) of each sample was diluted to 0.05, and cultures (100 µL) were added to 96 well plates (Thermo Scientific™ Nunc™ Edge™ 96-Well, Nunclon Delta-Treated, Flat-Bottom Microplate). OD was measured every 10 minutes for 20 hours at 37°C in a TECAN SPARK plate reader using a humidity cassette containing water to minimize evaporation. Growth curves were fit to the Gompertz equation using a custom R script with the nls function in the R stats package, version 3.6.2^55^.

### Library selection

The naive library (90 ng) was mixed with pLW002 (5 ng) and pLW003 (5 ng), the mixture was transformed into *E. coli* EW11 using electroporation, and cells were plated onto LB-agar medium containing chloramphenicol (34 µg/mL). After overnight growth, >10,000 CFU were observed, which was predicted to sample >99.9% coverage of the possible variants assuming sampling with replacement^54^. Colonies were harvested, plasmid DNA was purified to obtain *naive* library samples for sequencing, and the remaining slurry was used to inoculate three cultures (50 mL) containing selective (M9sa) and unselective (M9c) medium at an OD of 2x10^-6^. Cultures were grown for 12 hours at 37°C while shaking at 250 rpm, the final OD values were measured, and plasmids were purified to obtain selected and unselected library samples.

### Next Generation Sequencing

The full CysI gene and flanking region containing the barcode for the positive control was PCR amplified from the naïve, unselected, and selected samples. A commercial vendor (Azenta) fragmented the sequences, added indexes to each sample, and performed sequencing using Illumina MiSeq. This sequencing yielded >10M sequences for a mixture of each library. The forward and reverse sequencing data from paired reads were merged together using the Bbmerge.sh script^56^. A previously described script, dipseq, was used to identify and count each library variant under the different growth conditions^44^. Variant enrichment values were calculated using Rosace using default settings^57^.

### Modeling Fld binding to SiR-HP

The cryoelectron microscopy structure of the minimal SiR dimer was recently reported^29^, in which the Fld domain of SiR-FP is associated with the FNR domain. However, the Fld domain is dynamic and must dissociate from the FNR domain and shuttle electrons to SiR-HP^58–60^. To create a structural model for this conformational state, we modeled the Fld and SiR-HP interaction using AlphaFold-multimer by omitting the rest of the SiR-FP sequence. Target sequences were obtained from Uniprot for the *E. coli* SiR hemoprotein (P17846) and flavoprotein (P38038). We used ColabFold^61^ with multiple sequence alignments obtained from ColabFoldDB using the mmSEQs2 webserver^62,63^. Both paired and unpaired sequences were included in the alignment, and templates were automatically queried from pdb70^64^. Cofactors were placed in the AlphaFold models by backbone alignment with the crystal structures 6C3M and 6EFV. The predicted structure of the protein complex presented an average pLDDT of 95, pTM of 0.93, and interchain pTM of 0.82, suggesting high confidence. Residue pairs within each protein had low predicted aligned error (PAE), and despite high interchain PAEs, residues pairs at the interface were predicted with low PAE (Supplementary Fig. 18).

### Structural analysis

SiR is a large dynamic protein assembly made up of four SiR-HP and eight SiR-FP subunits, with multiple conformations important for function^58–60^. To evaluate how insertion tolerance relates to SiR-HP structural features, we evaluated how enrichment scores map onto the crystal structures of the free SiR-HP, PDB 6C3M^32^, the structure of the SiR-HP/FP dimer, PDB 9C91^29^, and the predicted Fld and SIR-HP complexes. Insertion proximity to cofactors and secondary structure analysis were performed using PDB 6C3M^32^. Trends in enrichment by secondary structure were determined by comparing the average enrichment across positions within loops, helices, and sheets to positions within the other two secondary structure types. To determine intramolecular contact density, residue-residue interactions were identified between the residue immediately preceding the peptide insertion site and other residues having heavy atoms within cutoffs of 4.5, 8, and 14 Å^65,66^. For the subunit interface in the SiR-HP/FP dimer and the modeled SiR-HP/Fld complex, the same method was used to identify intermolecular contacts.

### Electrochemical Measurements

Electrochemical measurements were performed in bioelectrochemical systems (BES) as described^67^. Colonies of *E. coli* EW11 transformed with constructs of interest were used to inoculate cultures of M9c (3 mL) containing 10 mM thiosulfate and antibiotic. After incubating at 30°C with shaking at 250 rpm for 12 hours, cultures were used to inoculate pre-cultures (100-fold dilution) of 50 mL M9c containing 10 mM thiosulfate, antibiotic, and either 10µM of 4-HT dissolved in ethanol or ethanol alone. These pre-cultures were grown to an OD of 0.6 at 30°C and 250 rpm. Bioreactors were set up by adding 120 mL of a modified M9sa (modifications included 0.2% glucose and exclusion of ferric citrate and magnesium sulfate) with 10mM thiosulfate; antibiotic was not included. Bioreactors, which were sparged with nitrogen gas prior, had carbon felt working electrodes poised at +300 mV against Ag/AgCl. After sparging, the precultures were washed three times with M9 salts before injecting into the bioreactors at an OD of 0.2. Bioreactors were incubated at 30°C while stirring at 300 rpm. Current density data was continuously collected via potentiostat from each bioreactor for ∼86-hours. Following setup of the bioreactors, SiR-HP activity was induced by injection of cells, thiosulfate, and either 4-HT (10 µM) or ethanol; this induction point is represented as time zero in all plots.

### Statistics

A two-component gaussian mixture model was fitted to the enrichment data using the Scikit-learn library version 1.2.2 for Python (jmlr.org/papers/v12/pedregosa11a.html). To quantify correlations between structural features and enrichment, Spearman correlations were calculated using the Scipy library version 1.10.1 for Python. The significance of Spearman correlations and differences in average enrichment for each secondary structure element were calculated by permutation tests with 100,000 resamples, also using Scipy. Paired one-tailed t-tests were conducted when quantifying the effect of 4-HT on domain variant growth. To determine the effect of estrogen antagonists on growth, a one-way ANOVA was performed followed by a post-hoc Dunnett’s test with DMSO serving as the control group. To determine the effect of 4-HT on the current density of the electrochemical measurements, one-tailed t-tests were performed.

## ACKNOWELDGEMENTS

We are grateful for support from the Office of Basic Energy Sciences of the U.S. Department of Energy grant DE-SC0014462 and National Science Foundation (NSF) grant 2223678. Research was sponsored by the Army Research Office and was accomplished under Grant Number W911NF-22-1-0239. The views and conclusions contained in this document are those of the authors and should not be interpreted as representing the official policies, either expressed or implied, of the Army Research Office or the U.S. Government. The U.S. Government is authorized to reproduce and distribute reprints for Government purposes notwithstanding any copyright notation herein. DM is currently supported by an NSF Graduate Research Fellowship.

## REFERENCES

1. Schwarzenbach, R. P. et al. The Challenge of Micropollutants in Aquatic Systems. Science 313, 1072–1077 (2006).

2. Vörösmarty, C. J. et al. Global threats to human water security and river biodiversity. Nature 467, 555–561 (2010).

3. Khalil, A. S. & Collins, J. J. Synthetic biology: applications come of age. Nat Rev Genet 11, 367–379 (2010).

4. Tsien, R. Y. THE GREEN FLUORESCENT PROTEIN. Annu. Rev. Biochem. 67, 509–544 (1998).

5. Contag, C. H. et al. Visualizing Gene Expression in Living Mammals Using a Bioluminescent Reporter. Photochem & Photobiology 66, 523–531 (1997).

6. Loper, J. E. & Henkels, M. D. Availability of iron to Pseudomonas fluorescens in rhizosphere and bulk soil evaluated with an ice nucleation reporter gene. Appl Environ Microbiol 63, 99–105 (1997).

7. Cheng, H.-Y. et al. Ratiometric Gas Reporting: A Nondisruptive Approach To Monitor Gene Expression in Soils. ACS Synth. Biol. 7, 903–911 (2018).

8. Li, S., Zuo, X., Carpenter, M. D., Verduzco, R. & Ajo-Franklin, C. M. Microbial bioelectronic sensors for environmental monitoring. Nat Rev Bioeng 3, 30–49 (2024).

9. Gralnick, J. A. & Bond, D. R. Electron Transfer Beyond the Outer Membrane: Putting Electrons to Rest. Annu. Rev. Microbiol. 77, 517–539 (2023).

10. Bird, L. J. et al. Engineering Wired Life: Synthetic Biology for Electroactive Bacteria. ACS Synth. Biol. 10, 2808–2823 (2021).

11. Golitsch, F., Bücking, C. & Gescher, J. Proof of principle for an engineered microbial biosensor based on Shewanella oneidensis outer membrane protein complexes. Biosensors and Bioelectronics 47, 285–291 (2013).

12. Webster, D. P. et al. An arsenic-specific biosensor with genetically engineered Shewanella oneidensis in a bioelectrochemical system. Biosensors and Bioelectronics 62, 320–324 (2014).

13. Ueki, T., Nevin, K. P., Woodard, T. L. & Lovley, D. R. Genetic switches and related tools for controlling gene expression and electrical outputs of *Geobacter sulfurreducens*. Journal of Industrial Microbiology and Biotechnology 43, 1561–1575 (2016).

14. West, E. A., Jain, A. & Gralnick, J. A. Engineering a Native Inducible Expression System in *Shewanella oneidensis* to Control Extracellular Electron Transfer. ACS Synth. Biol. 6, 1627–1634 (2017).

15. Dundas, C. M., Walker, D. J. F. & Keitz, B. K. Tuning Extracellular Electron Transfer by *Shewanella oneidensis* Using Transcriptional Logic Gates. ACS Synth. Biol. 9, 2301–2315 (2020).

16. Zhang, X. & Ajo-Franklin, C. Multichannel bioelectronic sensing using engineered *Escherichia coli*. Preprint at 10.1101/2023.09.30.560307 (2023).

17. Atkinson, J. T. et al. Real-time bioelectronic sensing of environmental contaminants. Nature 611, 548–553 (2022).

18. Cai, R. et al. Creation of a point-of-care therapeutics sensor using protein engineering, electrochemical sensing and electronic integration. Nat Commun 15, 1689 (2024).

19. Guo, Z. et al. Generalizable Protein Biosensors Based on Synthetic Switch Modules. J. Am. Chem. Soc. 141, 8128–8135 (2019).

20. Guo, Z. et al. Engineered PQQ-Glucose Dehydrogenase as a Universal Biosensor Platform. J. Am. Chem. Soc. 138, 10108–10111 (2016).

21. Mathai, J. C. et al. No facilitator required for membrane transport of hydrogen sulfide. Proc. Natl. Acad. Sci. U.S.A. 106, 16633–16638 (2009).

22. Crane, B. R., Siegel, L. M. & Getzoff, E. D. Sulfite Reductase Structure at 1.6 Å: Evolution and Catalysis for Reduction of Inorganic Anions. Science 270, 59–67 (1995).

23. Tavolieri, A. M. et al. NADPH-dependent sulfite reductase flavoprotein adopts an extended conformation unique to this diflavin reductase. Journal of Structural Biology 205, 170–179 (2019).

24. Barstow, B. et al. A synthetic system links FeFe-hydrogenases to essential E. coli sulfur metabolism. J Biol Eng 5, 7 (2011).

25. Coyote-Maestas, W., Nedrud, D., Okorafor, S., He, Y. & Schmidt, D. Targeted insertional mutagenesis libraries for deep domain insertion profiling. Nucleic Acids Res 48, e11 (2020).

26. Gonzalez, C. E., Roberts, P. & Ostermeier, M. Fitness Effects of Single Amino Acid Insertions and Deletions in TEM-1 β-Lactamase. Journal of Molecular Biology 431, 2320–2330 (2019).

27. Truong, A., Myerscough, D., Campbell, I., Atkinson, J. & Silberg, J. J. A cellular selection identifies elongated flavodoxins that support electron transfer to sulfite reductase. Protein Science 32, e4746 (2023).

28. Campbell, I. J. et al. Determinants of Multiheme Cytochrome Extracellular Electron Transfer Uncovered by Systematic Peptide Insertion. Biochemistry 61, 1337–1350 (2022).

29. Esfahani, B. G. et al. Structure of dimerized assimilatory NADPH-dependent sulfite reductase reveals the minimal interface for diflavin reductase binding. Preprint at 10.1101/2024.06.14.599029 (2024).

30. Jumper, J. et al. Highly accurate protein structure prediction with AlphaFold. Nature 596, 583–589 (2021).

31. Kim, J. Y., Nakayama, M., Toyota, H., Kurisu, G. & Hase, T. Structural and mutational studies of an electron transfer complex of maize sulfite reductase and ferredoxin. J Biochem 160, 101–109 (2016).

32. Cepeda, M. R., McGarry, L., Pennington, J. M., Krzystek, J. & Stroupe, M. E. The role of extended Fe4S4 cluster ligands in mediating sulfite reductase hemoprotein activity. Biochimica et Biophysica Acta (BBA) - Proteins and Proteomics 1866, 933– 940 (2018).

33. Shiau, A. K. et al. The Structural Basis of Estrogen Receptor/Coactivator Recognition and the Antagonism of This Interaction by Tamoxifen. Cell 95, 927–937 (1998).

34. Paige, L. A. et al. Estrogen receptor (ER) modulators each induce distinct conformational changes in ER α and ER β. Proc. Natl. Acad. Sci. U.S.A. 96, 3999– 4004 (1999).

35. Wu, B., Atkinson, J. T., Kahanda, D., Bennett, G. N. & Silberg, J. J. Combinatorial design of chemical-dependent protein switches for controlling intracellular electron transfer. AIChE Journal 66, e16796 (2020).

36. Atkinson, J. T. et al. Metalloprotein switches that display chemical-dependent electron transfer in cells. Nat Chem Biol 15, 189–195 (2019).

37. Truong, A. & Silberg, J. J. Regulating ferredoxin electron transfer using nanobody and antigen interactions. *RSC Chem*. Biol. (2025) doi:10.1039/D4CB00257A.

38. Atkinson, J. T. et al. Real-time bioelectronic sensing of environmental contaminants. Nature 611, 548–553 (2022).

39. Sekhon, H., Ha, J.-H. & Loh, S. N. Enhancing response of a protein conformational switch by using two disordered ligand binding domains. Front. Mol. Biosci. 10, 1114756 (2023).

40. Claus, M. T., Zocher, G. E., Maier, T. H. P. & Schulz, G. E. Structure of the *O* - Acetylserine Sulfhydrylase Isoenzyme CysM from *Escherichia coli*,. Biochemistry 44, 8620–8626 (2005).

41. Hryniewicz, M., Sirko, A., Pałucha, A., Böck, A. & Hulanicka, D. Sulfate and thiosulfate transport in Escherichia coli K-12: identification of a gene encoding a novel protein involved in thiosulfate binding. J Bacteriol 172, 3358–3366 (1990).

42. Brown, M. D., Hall, J. R. & Schoenfisch, M. H. A direct and selective electrochemical hydrogen sulfide sensor. Analytica Chimica Acta 1045, 67–76 (2019).

43. Nadler, D. C., Morgan, S.-A., Flamholz, A., Kortright, K. E. & Savage, D. F. Rapid construction of metabolite biosensors using domain-insertion profiling. Nat Commun 7, 12266 (2016).

44. Oakes, B. L. et al. Profiling of engineering hotspots identifies an allosteric CRISPR- Cas9 switch. Nat Biotechnol 34, 646–651 (2016).

45. Coyote-Maestas, W., He, Y., Myers, C. L. & Schmidt, D. Domain insertion permissibility-guided engineering of allostery in ion channels. Nat Commun 10, 290 (2019).

46. Wolf, B. et al. Rational engineering of allosteric protein switches by *in silico* prediction of domain insertion sites. Preprint at 10.1101/2024.12.04.626757 (2024).

47. VanArsdale, E. et al. A Coculture Based Tyrosine-Tyrosinase Electrochemical Gene Circuit for Connecting Cellular Communication with Electronic Networks. ACS Synth. Biol. 9, 1117–1128 (2020).

48. 48. Ajo-Franklin, C., Zhang, X. & Charrier, M. Multichannel bioelectronic sensing using engineered Escherichia coli. Preprint at 10.21203/rs.3.rs-4853950/v1 (2024).

49. Li, S., Tavares, C. D. G., Tolar, J. G. & Ajo-Franklin, C. M. Selective bioelectronic sensing of quinone pharmaceuticals using extracellular electron transfer in *Lactiplantibacillus plantarum*. Preprint at 10.1101/2023.03.23.533500 (2023).

50. Tschirhart, T. et al. Electronic control of gene expression and cell behaviour in Escherichia coli through redox signalling. Nat Commun 8, 14030 (2017).

51. Smith, K. W. & Stroupe, M. E. Mutational Analysis of Sulfite Reductase Hemoprotein Reveals the Mechanism for Coordinated Electron and Proton Transfer. Biochemistry 51, 9857–9868 (2012).

52. Morley, K. L. & Kazlauskas, R. J. Improving enzyme properties: when are closer mutations better? Trends in Biotechnology 23, 231–237 (2005).

53. Engler, C., Kandzia, R. & Marillonnet, S. A One Pot, One Step, Precision Cloning Method with High Throughput Capability. PLoS ONE 3, e3647 (2008).

54. Bosley, A. D. & Ostermeier, M. Mathematical expressions useful in the construction, description and evaluation of protein libraries. Biomolecular Engineering 22, 57–61 (2005).

55. Zwietering, M. H., Jongenburger, I., Rombouts, F. M. & Van ’T Riet, K. Modeling of the Bacterial Growth Curve. Appl Environ Microbiol 56, 1875–1881 (1990).

56. Bushnell, B., Rood, J. & Singer, E. BBMerge – Accurate paired shotgun read merging via overlap. PLoS ONE 12, e0185056 (2017).

57. Rao, J. et al. Rosace: a robust deep mutational scanning analysis framework employing position and mean-variance shrinkage. Genome Biol 25, 138 (2024).

58. Murray, D. T., Weiss, K. L., Stanley, C. B., Nagy, G. & Stroupe, M. E. Small-angle neutron scattering solution structures of NADPH-dependent sulfite reductase. Journal of Structural Biology 213, 107724 (2021).

59. Murray, D. T. et al. Neutron scattering maps the higher-order assembly of NADPH-dependent assimilatory sulfite reductase. Biophysical Journal 121, 1799–1812 (2022).

60. Walia, N. et al. Domain crossover in the reductase subunit of NADPH-dependent assimilatory sulfite reductase. Journal of Structural Biology 215, 108028 (2023).

61. Mirdita, M. et al. ColabFold: making protein folding accessible to all. Nat Methods 19, 679–682 (2022).

62. Steinegger, M. & Söding, J. MMseqs2 enables sensitive protein sequence searching for the analysis of massive data sets. Nat Biotechnol 35, 1026–1028 (2017).

63. Mirdita, M., Steinegger, M. & Söding, J. MMseqs2 desktop and local web server app for fast, interactive sequence searches. Bioinformatics 35, 2856–2858 (2019).

64. Berman, H. M. The Protein Data Bank. Nucleic Acids Research 28, 235–242 (2000).

65. Kozma, D., Simon, I. & Tusnady, G. E. CMWeb: an interactive on-line tool for analysing residue-residue contacts and contact prediction methods. Nucleic Acids Research 40, W329–W333 (2012).

66. Nagarajan, R. et al. PDBparam: Online Resource for Computing Structural Parameters of Proteins. Bioinform Biol Insights 10, BBI.S38423 (2016).

67. Alba, R. A. C., Li, S., Kundu, B. B., Ajo-Franklin, C. M. & Cai, R. Characterizing Mediated Extracellular Electron Transfer in Lactic Acid Bacteria with a Three-electrode, Two-chamber Bioelectrochemical System. JoVE 67204 (2024) doi:10.3791/67204.

