## Supporting Information for "Using domain insertion to create sulfite reductases that present chemical-dependent activities"

Author affiliations:

### TABLE OF CONTENTS


| Item | Title | Page |
|---|---|---|
| Fig. S1 | Growth complementation of *E. coli* EW11 by control plasmids | 3 |
| Fig. S2 | Library construction workflow and tile sequencing | 4 |
| Fig. S3 | Library analysis workflow and controls | 5 |
| Fig. S4 | NGS analysis of each naïve library replicate | 6 |
| Fig. S5 | NGS analysis of each unselected library replicate | 7 |
| Fig. S6 | NGS analysis of each selected library replicate | 8 |
| Fig. S7 | Average variant counts and CV for NGS | 9 |
| Fig. S8 | Peptide insertions are tolerated most in loops | 10 |
| Fig. S9 | Insertion tolerance and intermolecular interactions | 11 |
| Fig. S10 | Interface of the modeled Fld domain and SiR-HP complex | 12 |
| Fig. S11 | Comparison of enrichment values with prior HD-exchange data | 13 |
| Fig. S12 | Effect of 4-HT on domain insertion variant growth complementation | 14 |
| Fig. S13 | Maximum OD observed with growth complementation ±4-HT | 15 |
| Fig. S14 | Domain insertion variants analyzed under non-selective conditions | 16 |
| Fig. S15 | Domain insertion variant complementation ±ER-LBD ligands | 17 |
| Fig. S16 | Individual bioreactor measurements | 18 |
| Fig. S17 | Biomass measurements of individual replicates in bioreactors | 19 |
| Fig. S18 | Predicted alignment error for AlphaFold Fld/SiR-HP model | 20 |
| Table S1 | Peptide insertion variants targeted for domain insertion | 21 |
| Table S2 | Plasmids used in this study | 22 |

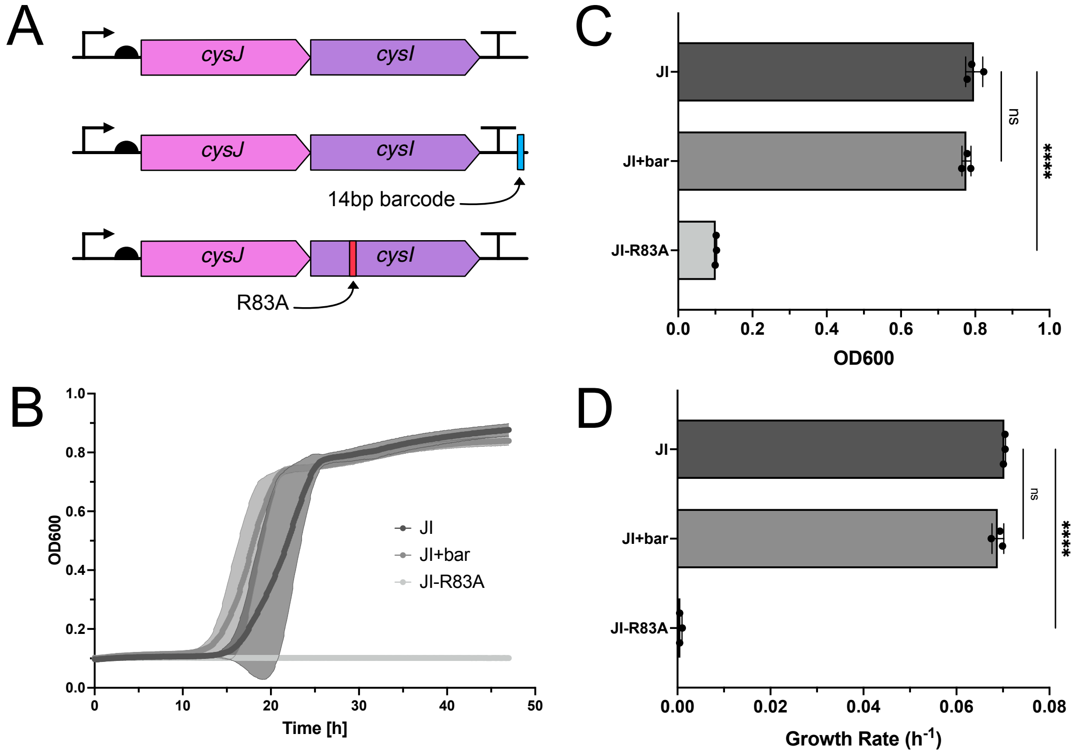


**Supplementary Figure 1. Growth complementation of *E. coli* EW11 by control plasmids.** (**A**) SIR expression plasmids used for this study. CysJI (JI) is the parent vector used to create the library. The CysJI plasmid having a 14 base pair barcode (JI+bar) was used as a positive control vector in library selections. The CysJI plasmid having the R83A mutation (JI-R83A) represents the negative control, which has a point mutation within the SiR-HP active site. (**B**) Growth of *E. coli* EW11 transformed with each construct in selective media at 37°C with shaking at 250 rpm. Lines represent the average from three biological replicates, while shaded regions indicate ±1$\sigma$. (**C**) $OD_{600}$ values from growth curves at 30 hours. The growth of JI+bar does not differ from JI (ns, p = 0.2, Dunnett's test), while the complementation with JI-R83A is significantly lower than the JI parent vector (****, p = 8e-07). Points represent individual biological replicates, bars represent the average, and error is shown ±1 standard deviation. (**D**) The growth rate of the JI-R83A mutant is significantly lower than JI (****, p = 6e-10), while the growth rate of JI+bar is not significantly different than JI (ns, p = 0.1).

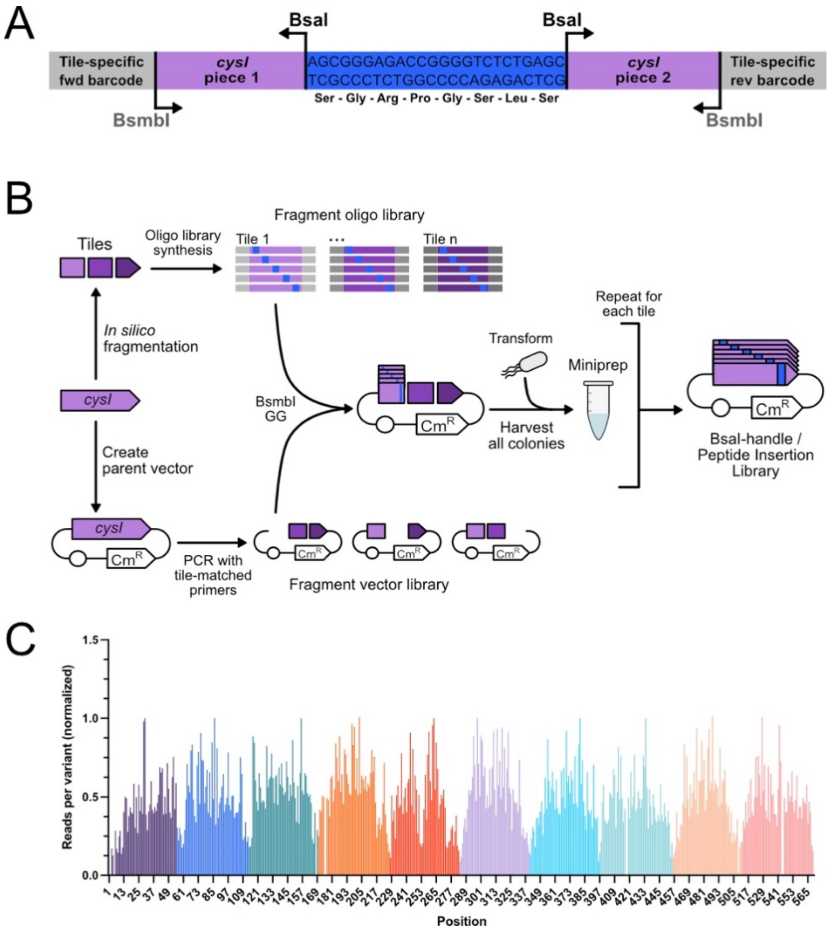


**Supplementary Figure 2. Library construction workflow and tile sequencing.** (**A**) Schematic of a representative variant, showing the tile-specific barcodes on either end, the BsmbI cut sites that flank the *cysI* sequence, and the octapeptide insertion occurring at some locus within the protein coding sequence. (**B**) Workflow for library construction. The SiR-HP gene is first fragmented into ten tiles in silico to design the library. Each tile gene is synthesized as an oligonucleotide pool with insertions (blue) at every possible location. Each oligonucleotide pool is cloned using Golden Gate assembly into the parent vector (JI in Supplemental Figure 1) to yield a plasmid ensemble for each tile. (**C**) Sequence diversity for the plasmids encoding each tile are shown in different colors. AmpliconEZ sequencing was performed individually with each tile to assess coverage, and the data from each reaction was merged to show the relative coverage of insertion variants. A single sequencing run was performed with each tile sample.

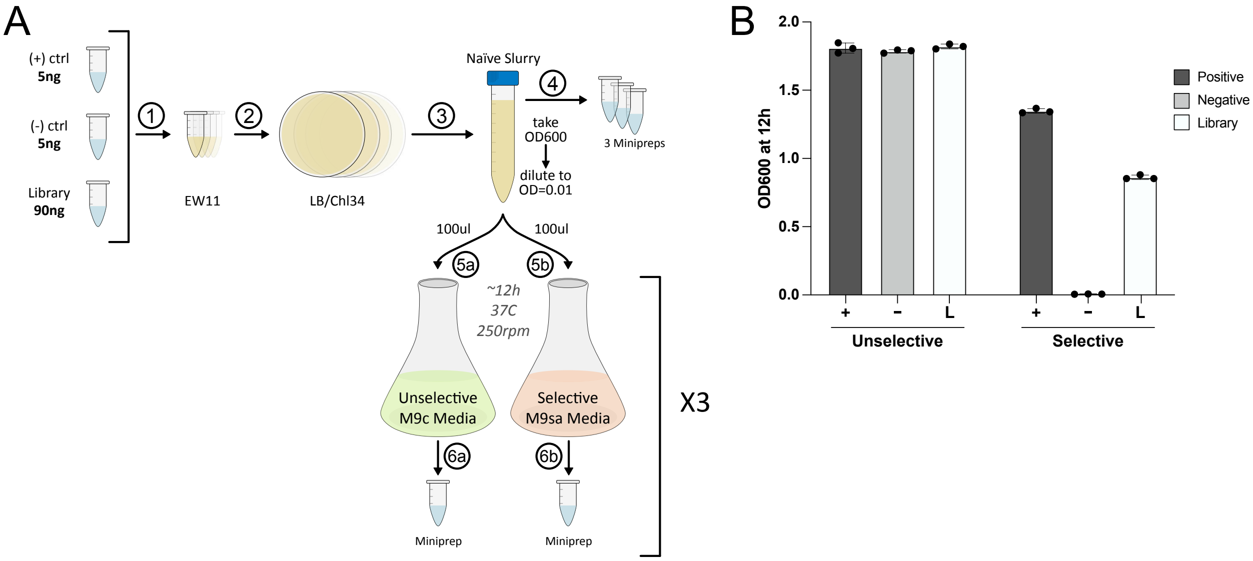


**Supplementary Figure 3. Library selection workflow and controls.** (**A**) Workflow for the library selection process: (1) after pooling tiles to create the full library, this mixture was combined with controls (JI and JI-R83A) and transformed into *E. coli* EW11 via electroporation, (2) transformants were grown on LB-agar medium at 37°C overnight, (3) colonies were harvested and pooled, (4) DNA was purified from three aliquots of the slurry to obtain the naïve library replicates, (5) an aliquot of the slurry was diluted to an $OD_{600}$ of 0.01 and used to inoculate three cultures containing either unselective or selective medium, and (6) cells were grown for 12 hours at 37°C with shaking at 250 rpm. To obtain the selective and unselective samples, DNA plasmids were purified from each culture following incubation. (**B**) $OD_{600}$ values for the positive (JI-bar) and negative controls (JI-R83A), and library following 12 hours of growth in selective and unselective medium. Biological replicates are shown as points (n = 3), bars represent the average, and error is shown as $\pm 1\sigma$. In the unselective condition, the $OD_{600}$ at 12 hours of the positive control was not significantly different from the negative control (ns, p = 0.4, Dunnet's test) or the library (ns, p = 0.6). In the selective condition, the $OD_{600}$ at 12 hours of the positive control was significantly higher than the negative control (p = 0.00005) and library (p = 0.000004).

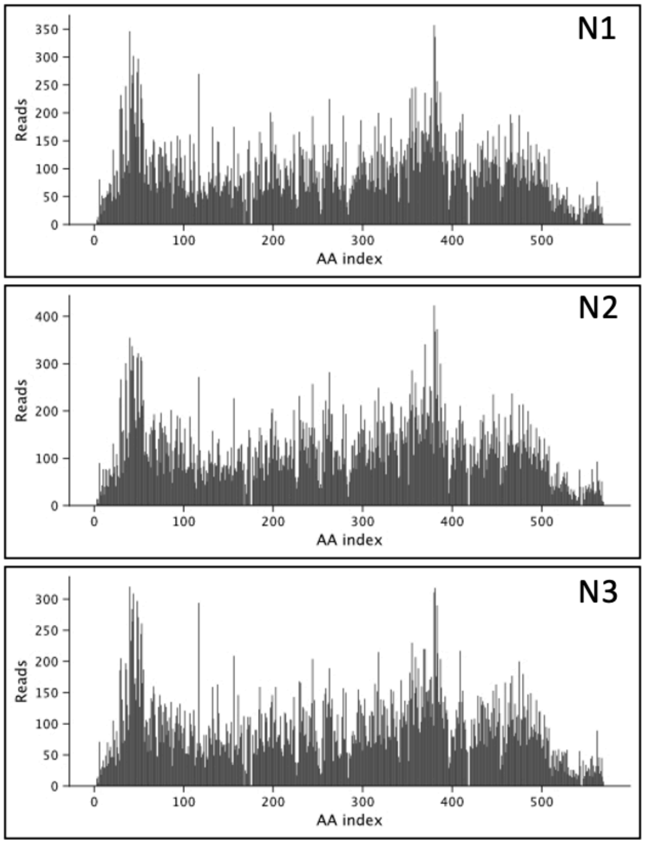


**Supplementary Figure 4. NGS analysis of each naïve library replicate.** N1, N2, and N3 show the number of unique reads observed for each peptide insertion variant from the three different biological replicates.

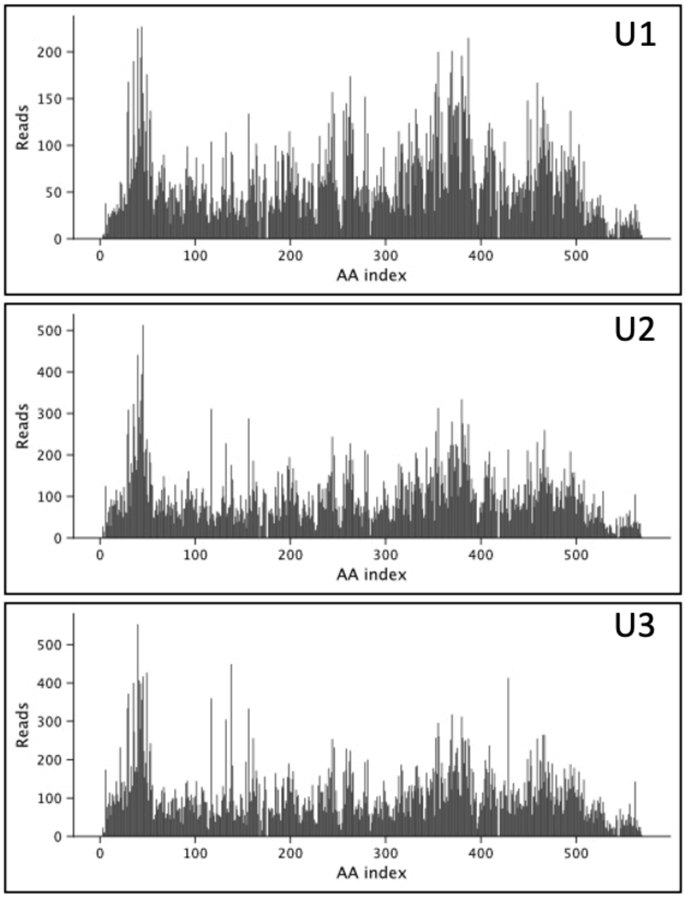


**Supplementary Figure 5. NGS analysis of each unselected library replicate.** U1, U2, and U3 represent the number of unique reads observed for each peptide insertion variant for the three different biological replicates from growth in the unselective medium.

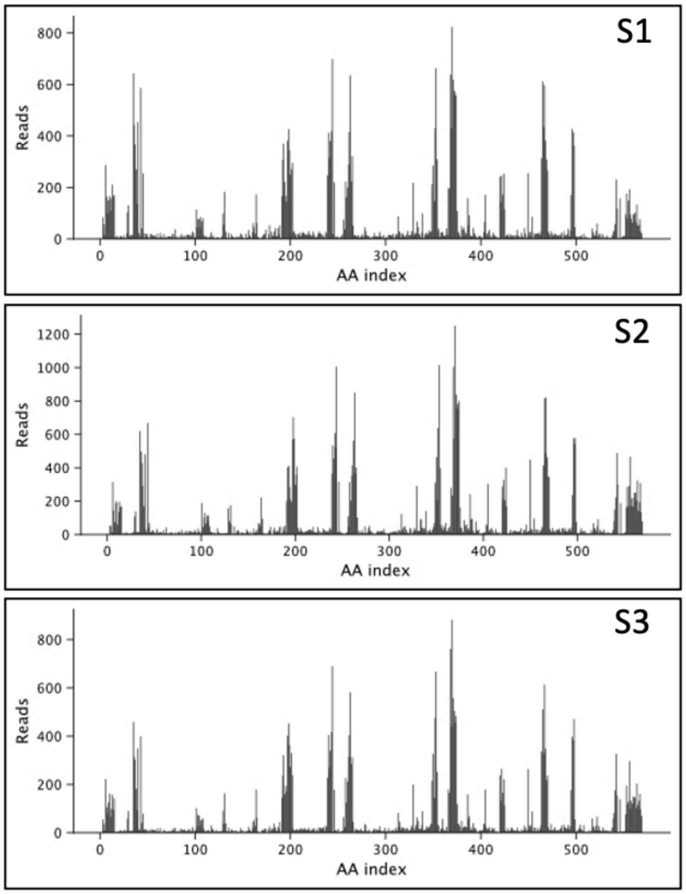


**Supplementary Figure 6. NGS analysis of each selected library replicate.** S1, S2, and S3 represent the number of unique reads observed for each peptide insertion variant from the three different biological replicates from growth in the selective medium.

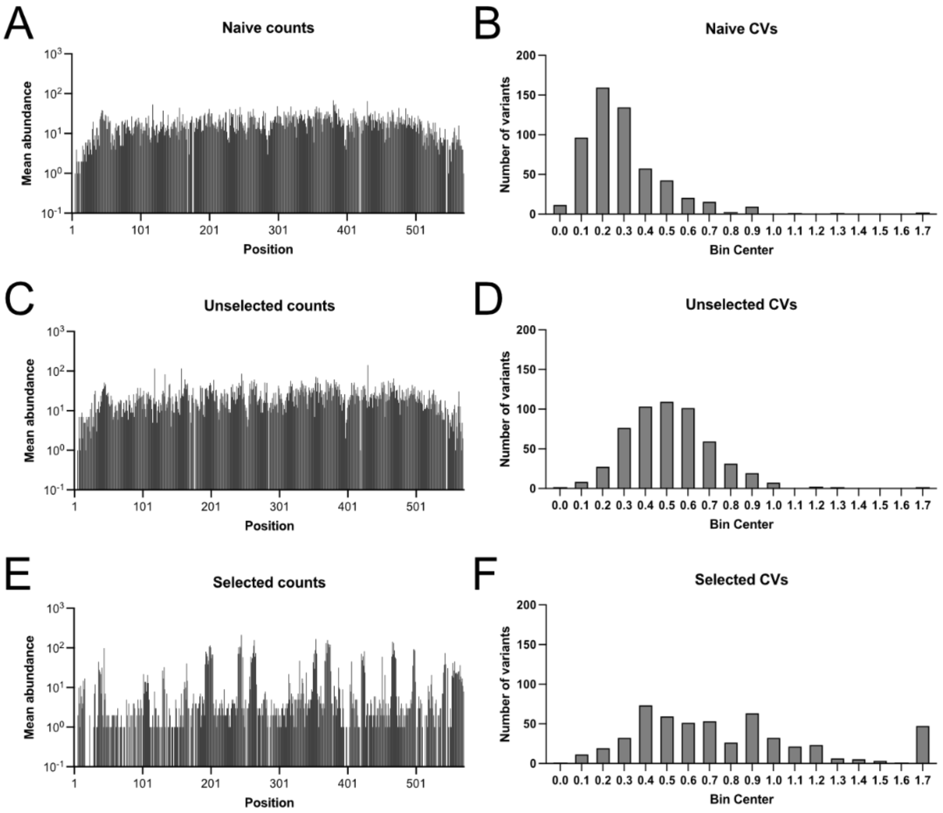


**Supplementary Figure 7. Average variant counts and CV for NGS.** The average sequence abundances for each insertion variant are shown for the (**A**) naïve, (**C**) unselected, and (**E**) selected samples. Histograms are shown for the CVs of read abundances for the (**B**) naive, (**D**) unselected, and (**F**) selected. The data shown was calculated using the data shown in Supplementary Figures 4, 5, and 6.

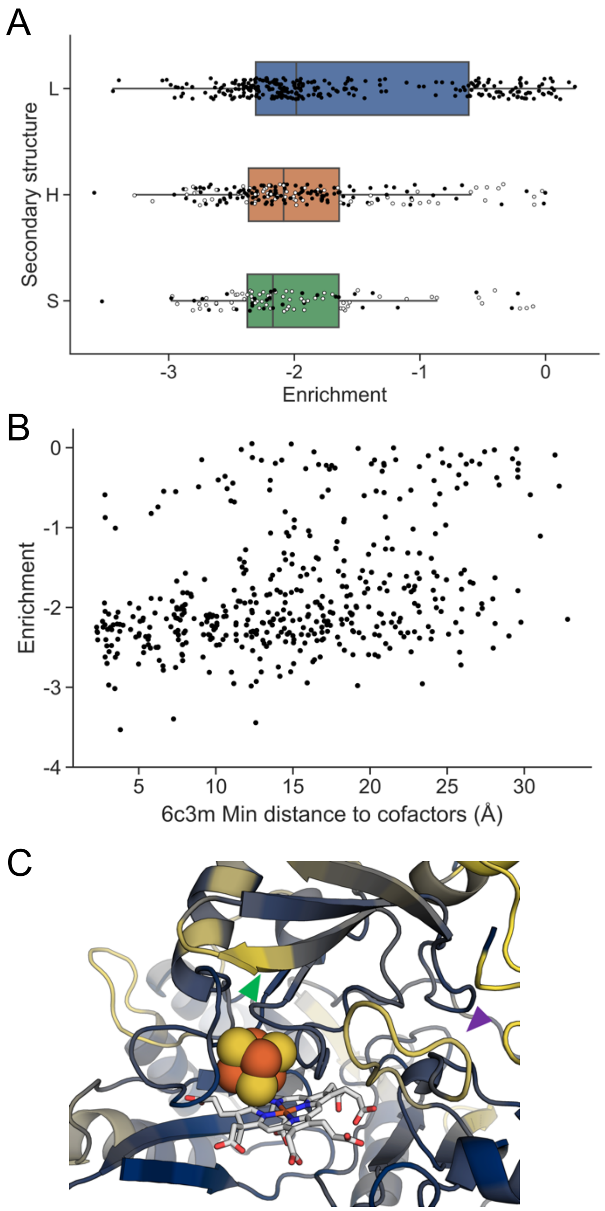


**Supplementary Figure 8. Peptide insertions are tolerated most in loops.** (**A**) Dots show that the average enrichment of insertions within loops (L, blue) is significantly higher than the other secondary structures ($p = 2.0x10^{-5}$, permutation test with 100000 resamples), including helices (H, orange) and sheets (S, green). Open circles are positions adjacent to a loop. (**B**) Enrichment versus minimum distance to a heavy atom in either cofactor shows a bimodality and higher enrichment being more common as positions with longer minimum distance to cofactor. (**C**) Zoom-in of the cofactors shows that there are two proximal loops that show high enrichment (yellow). DV496 bears a ER-LBD insertion within the beta strand above the Fe-S cluster (green triangle), and DV264 bears a ER-LBD insertion within the loop to the right of the cofactors (purple triangle).

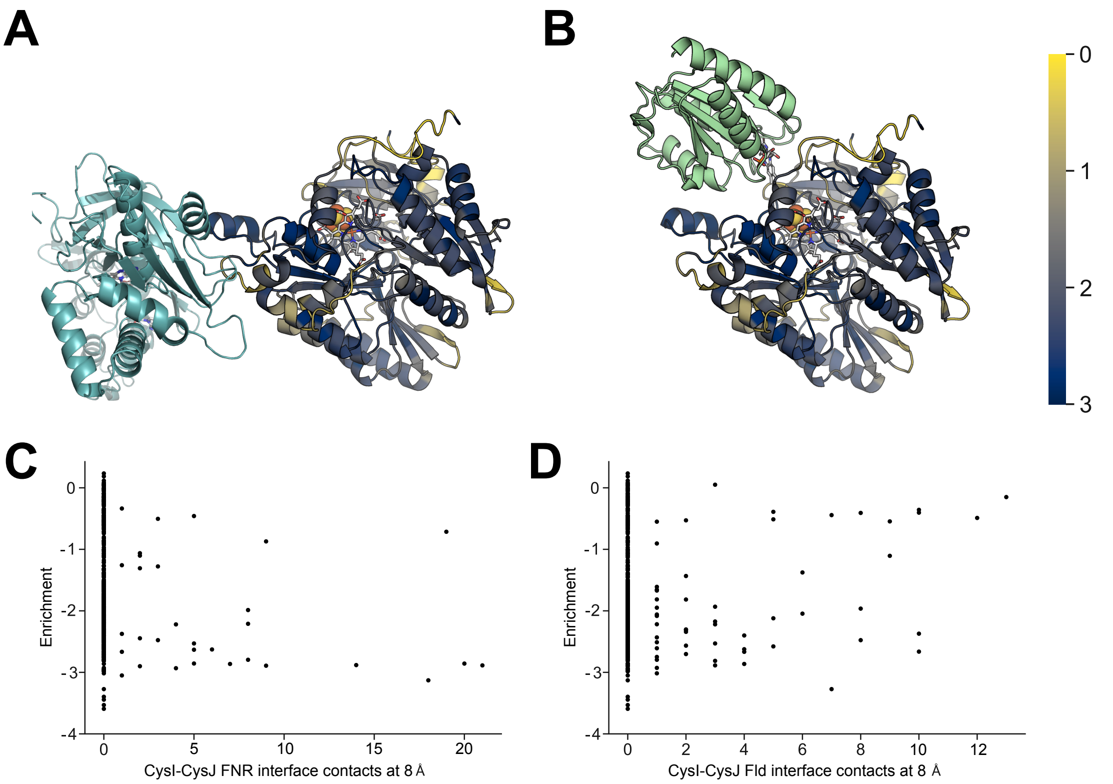


**Supplementary Figure 9. Peptide insertion tolerance and intermolecular interaction.** (**A**) The structure of the SIR-FP (cyan) and SIR-HP complex (PDB 9c91) determined using cryoelectron microscopy. SiR-HP is colored by the enrichment score obtained from library selections. (**B**) A model of a complex between the SiR-HP and the Fld domain of SiR-FP (green). In this structure, the flavin mononucleotide in the Fld domain is 8.8Å from the 4Fe-4S cofactor in SiR-HP. (**C**) At each site of peptide insertion in SiR-HP, the enrichment value is compared with the number of intermolecular residue-residue contacts (≤8 Å) made by residues preceding each insertion site in SiR-FP and residues in SIR-FP within the structure shown in panel A. (**D**) A comparison of enrichment values for SIR-HP insertions and the number of intermolecular contacts made between the residue preceding the insertion site in SiR-HP and the Fld domain of SiR-FP.

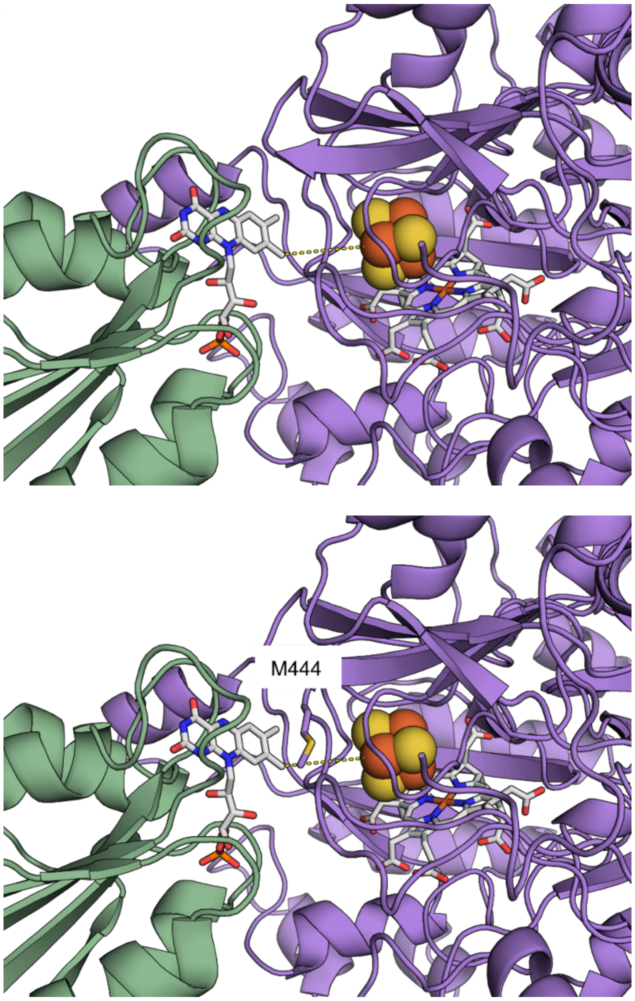


**Supplementary Figure 10. Interface of the modeled Fld domain and SiR-HP complex.** (*top*) Distance between the FMN and 4Fe-4S. A view of the predicted Fld/SiR-HP complex highlighting the shortest inter-atomic distance between the cofactors (dashed yellow line), which is 8.8 Å. SiR-HP is shown in purple and the SiR-FP Fld domain is shown in green. (*bottom*) The AlphaFold-predicted binding mode for SiR-HP and the SiR-FP Fld domain is consistent with reported mutational and biochemical data. A view of the predicted Fld/SiR-HP complex showing methionine 444, a residue thought to be involved in Fld domain binding and/or electron transfer to SiR-HP[1]. In the predicted complex, M444 is directly between the flavin and 4Fe-4S cofactors.

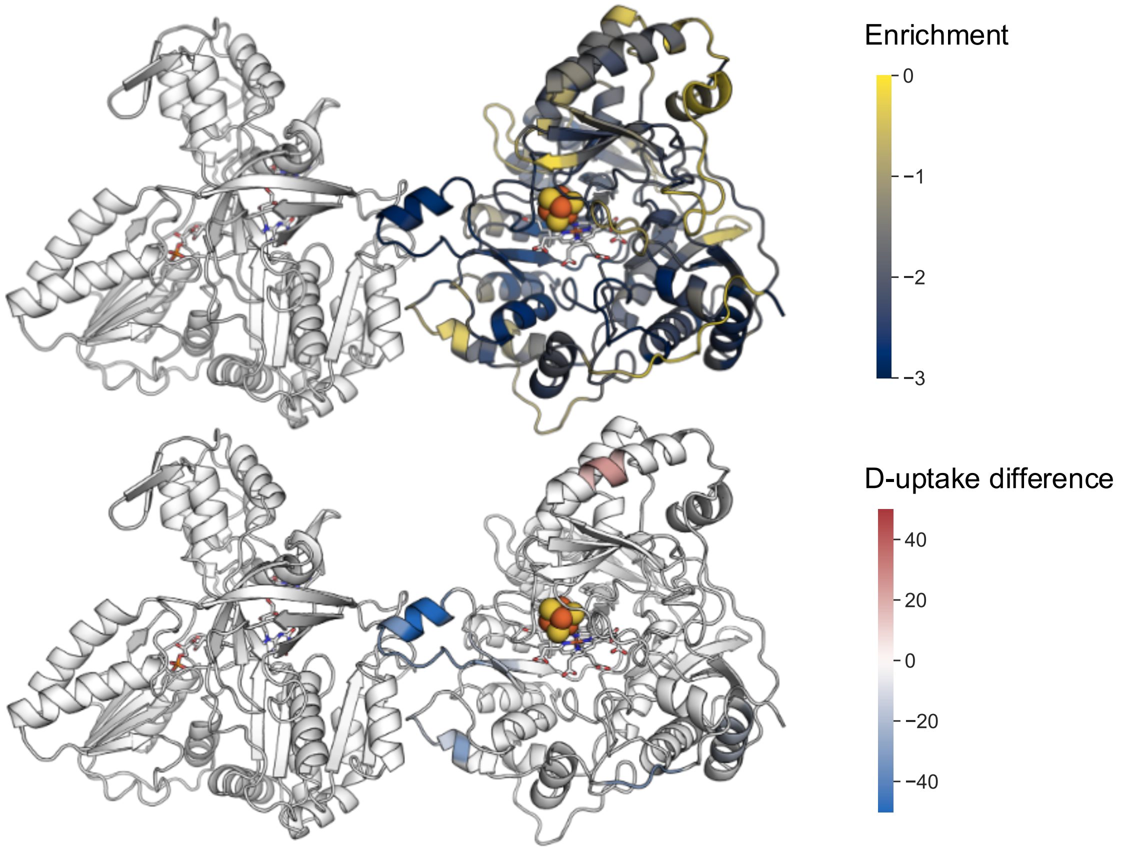


**Supplementary Figure 11. Comparison of enrichment values with prior HD-exchange data.** A view of the cryoelectron microscopy structure of the SiR dimer[2] shaded by: (*top*) enrichment and (*bottom*) HD-exchange data from Askenasy et al.[3]. SiR-HP residues with lower deuterium uptake in the SiR dodecamer compared to the monomer are shown in dark blue, while residues with greater uptake are shown in pink. Together, this data shows that the residues implicated as buried at the SiR-FP/HP interface by HD-exchange experiments are not all equally intolerant to peptide insertion.

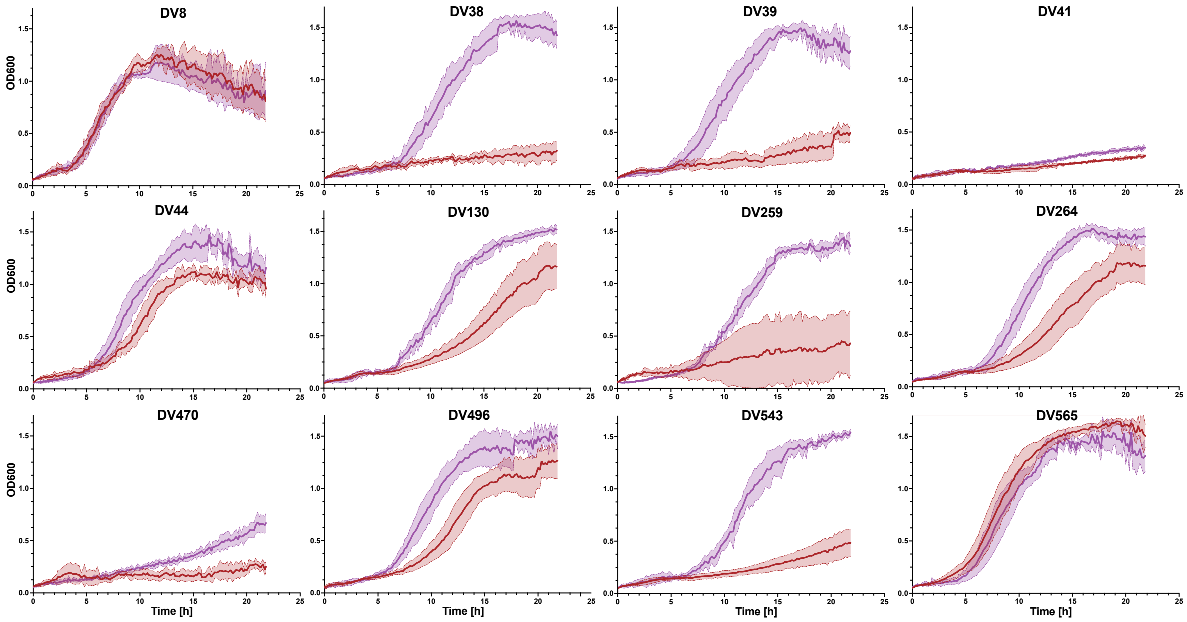


**Supplementary Figure 12. Effect of 4-HT on domain insertion variant growth complementation.** Growth complementation was measured in selective medium at 37°C while shaking at 250 rpm. The red traces indicate the average growth with addition of DMSO only, while the purple traces indicate the average growth when 4-HT was added to the growth medium. The shaded regions represent ±1 standard deviation from the six biological replicates.

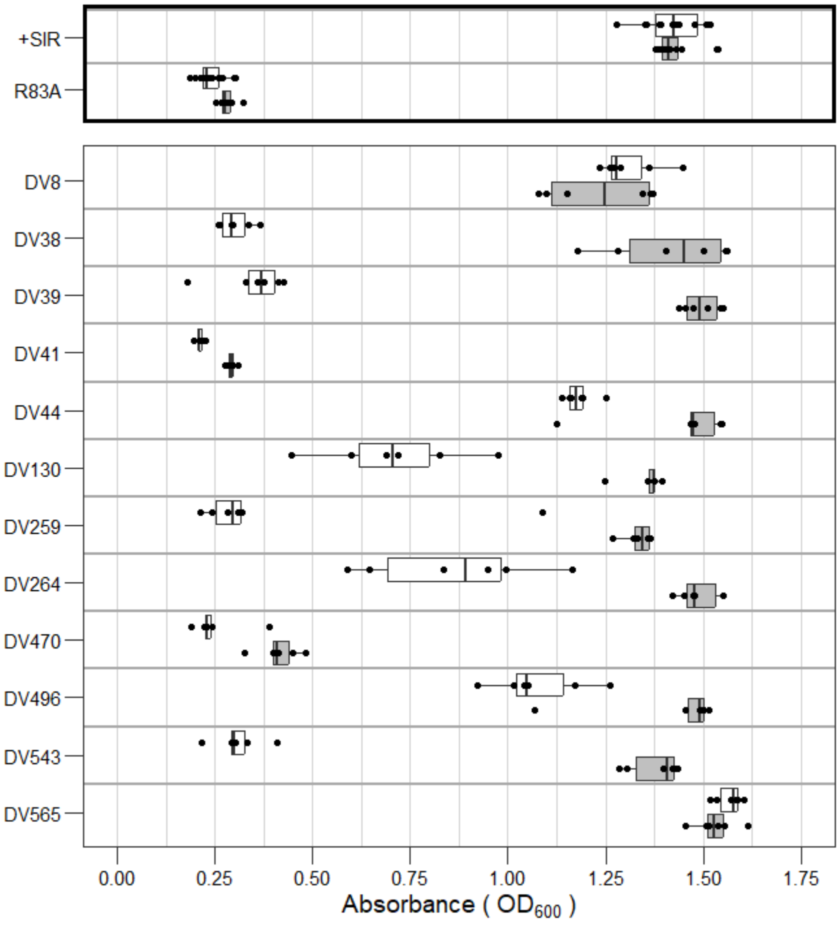


**Supplementary Figure 13. Maximum OD observed with growth complementation ±4-HT. (*top*)** Effect of 10 µM 4-HT (gray) and the DMSO vehicle (white) on the growth of *E. coli* EW11 expressing SiR-HP and SiR-FP (+SiR) or a R83A mutant of SiR-HP and SiR-FP (R83A) after 16 hours in selective medium. **(*bottom*)** Effect of 4-HT and DMSO on the growth of *E. coli* EW11 expressing SiR-FP and SiR-HP having the ER-LBD inserted in twelve locations. For all experiments, six biological replicates were performed (dots). Within each box, the center line represents the $50^{th}$ percentile, while ends of the box represent the $25^{th}$ and $75^{th}$ percentiles, respectively. Outside the box, the line tips represent the minima and maxima. Dots outside of the lines correspond to outliers. The growth of SiR-HP and domain variants +SiR, DV8, and DV565 were not enhanced by 4-HT ($p > 0.05$, one-tailed paired t-test), whereas variant DV38, DV39, DV41, DV44, DV130, DV259, DV264, DV470, DV496, and DV543 presented a significant increase in OD (DV38, p = 6e-06; DV39, p = 1e-06; DV41, p = 2e-06; DV44, p = 0.01; DV130, 0.0003; DV259, p = 0.0005; DV264, p = 0.0007; DV470, p = 0.002; DV496, p = 0.003; DV543, p = 8e-07, one-tailed paired t-test)**.** A small but significant increase in OD was observed with the R83A negative control p = 0.0007)

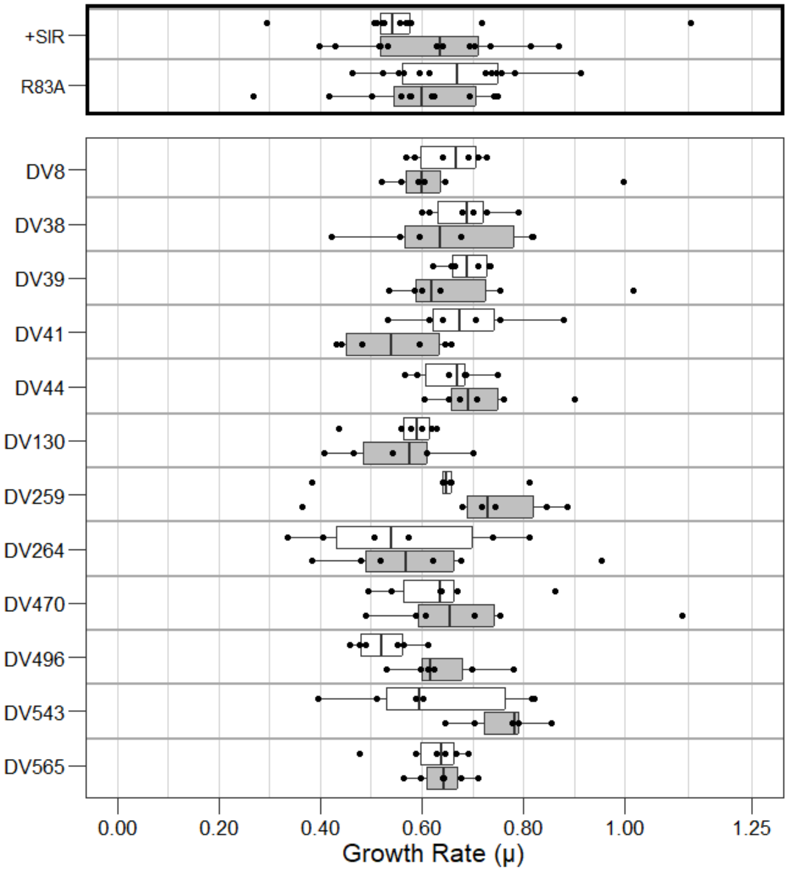


**Supplementary Figure 14. Domain insertion variants analyzed under non-selective conditions. (*top*)** Effect of 10 μM 4-HT (gray) and the DMSO vehicle (white) on the growth of *E. coli* EW11 expressing SiR-HP and SiR-FP (+SiR) or a R83A mutant of SiR-HP and SiR-FP (R83A). **(*bottom*)** Effect of 4-HT and DMSO on the growth of *E. coli* EW11 expressing SiR-FP and SiR-HP having the ER-LBD inserted at different locations. For all experiments, six biological replicates were performed (dots). Within each box, the center line represents the $50^{th}$ percentile, while ends of the box represent the $25^{th}$ and $75^{th}$ percentiles, respectively. Outside the box, the line tips represent the minima and maxima. Dots outside of the lines correspond to outliers. The growth of SiR-HP and domain variants 8, 38, 39, 41, 44, 130, 259, 264, 470, 543, and 565 were not enhanced by 4-HT ($p > 0.05$, one-tailed paired t-test), whereas domain variants 496 had significantly enhanced growth rate ($p = 0.039$, one-tailed paired t-test).

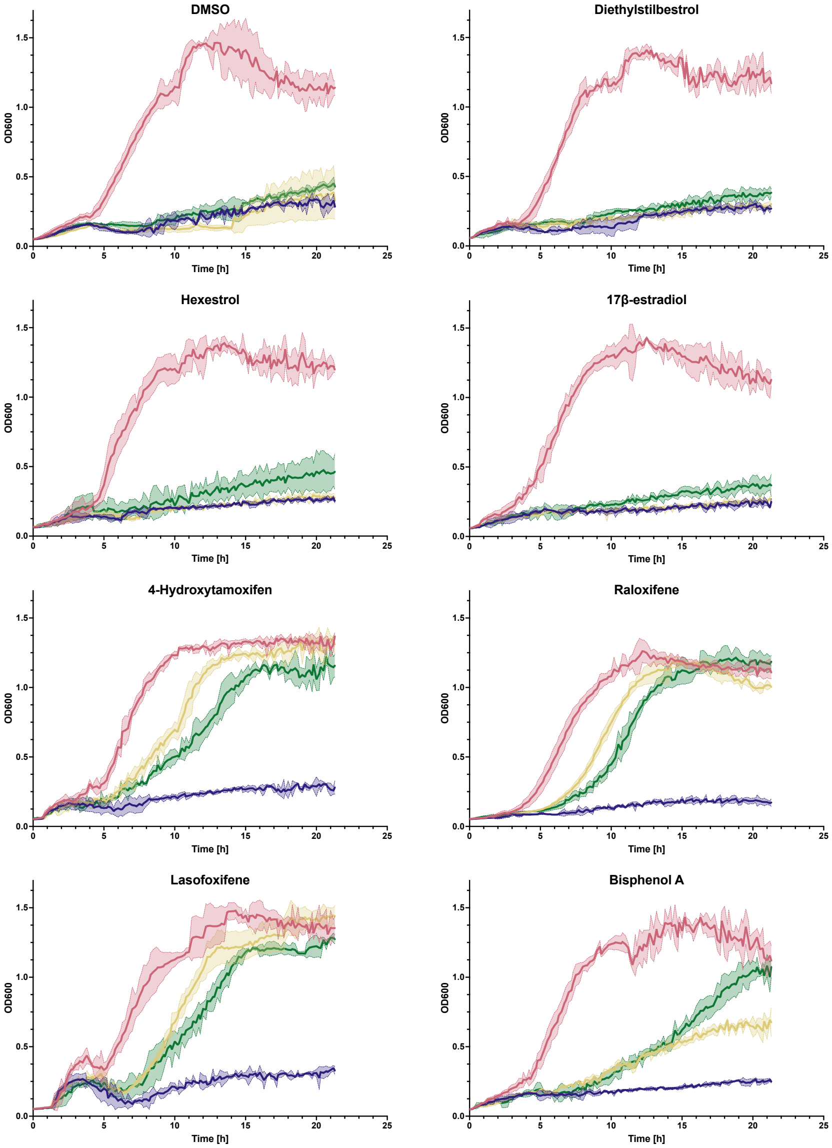


**Supplementary Figure 15. Domain insertion variant complementation ±ER-LBD ligands.** Growth complementation for each of the four constructs tested in selective medium containing eight different additions. The red traces represent the positive control (native SiR-HP), the blue traces represent for the negative control (R83A SiR-HP mutant), the green traces are for DV259, and the yellow traces are for DV39. Lines represent the average growth from three biological replicates at 37°C with shaking, while the shaded regions represent ±1 standard deviation.

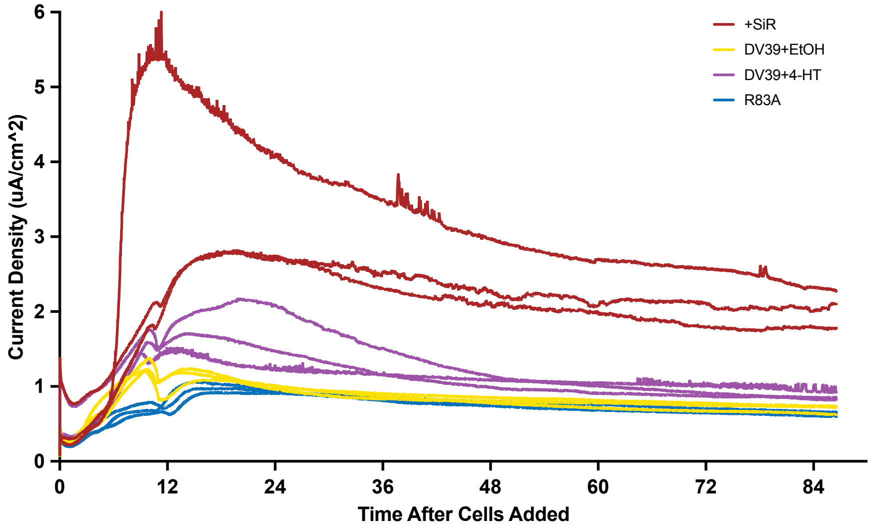


**Supplementary Figure 16. Individual bioreactor measurements.** Individual traces from the BESs. Red traces are the positive control with ethanol only, the blue traces are the inactive SiR-HP R83A mutant in ethanol, the purple traces are the DV39 switch with 4-HT in ethanol, and the yellow traces are DV39 switch with ethanol only. One of the positive control traces is much higher than the other two.

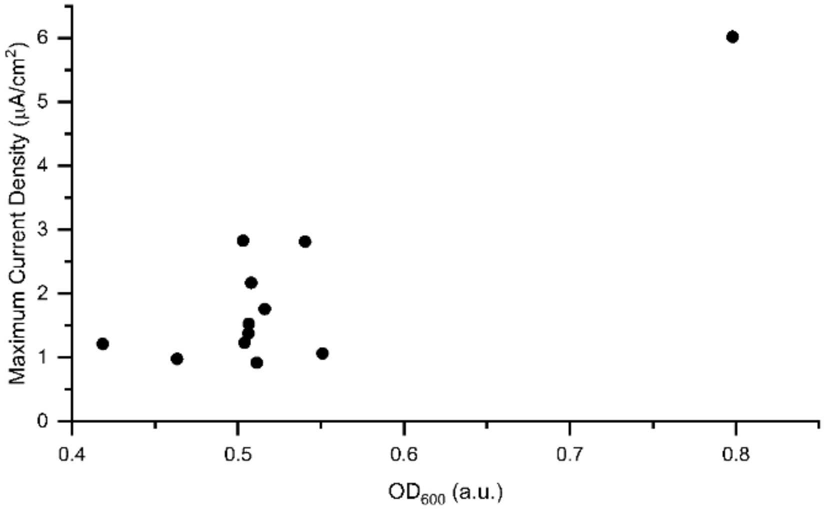


**Supplementary Figure 17. Biomass measurements of individual replicates in bioreactors.** Comparison of maximum current density with cell density, optical density at 600 nm ($OD_{600}$) for each individual sample. The outlier which shows high maximum current density also shows a much higher cell density than the other samples.

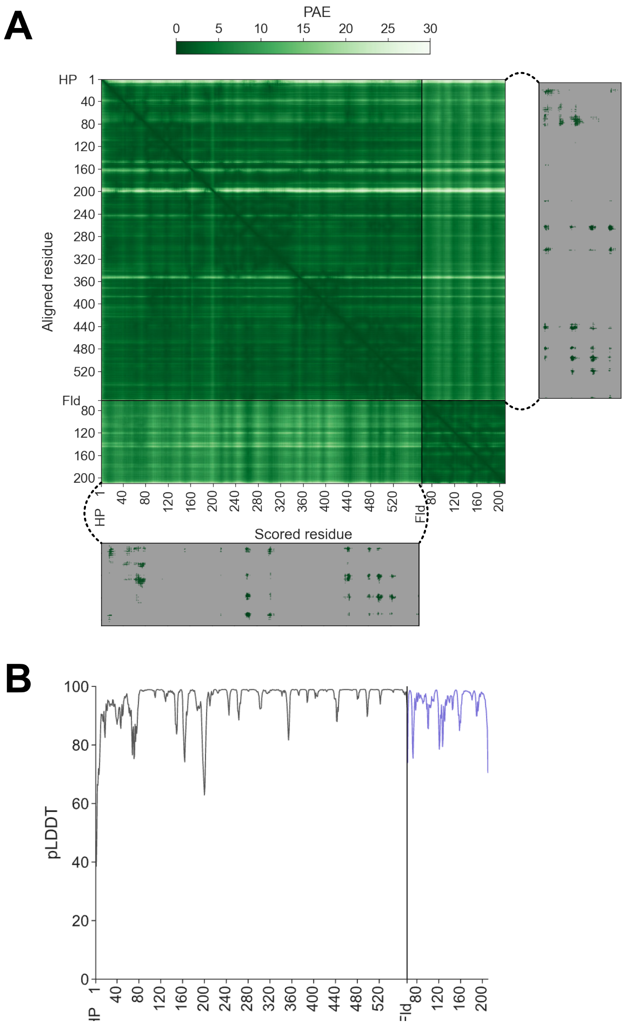


**Supplementary Figure 18. Predicted alignment error for AlphaFold Fld/SiR-HP model.** (**A**) Predicted aligned errors (PAEs) for the Fld/SiR-HP complex. Cutaways display PAEs for pairs of residues containing at least one heavy atom within 14 Å, with all others masked in gray, indicating that residue pairs at the Fld/SiR-HP interface are confidently predicted. (**B**) Plot of pLDDT for the predicted SiR-HP and Fld. The average pLDDT of SiR-HP is 95.8, and the average for the Fld domain is 94.1.

**Supplementary Table 1. Peptide insertion variants targeted for domain insertion.** List of the twelve domain inserted variants chosen to move forward with, grouped by their locality to interfaces and listed with their positions, enrichment scores, and number of contacts at the structural and dynamic interfaces. The dynamic interface represents the intermolecular contacts made by the SiR-FP Fld domain and SiR-HP, while the structural interface represents the intermolecular contacts observed in the cryoelectron microscopy structure of the SiR-FP/HP complex. The ER-LBD was inserted following the residue noted for each peptide insertion variant.

| Peptide insertion variant | Enrichment Score | Contacts at Structural Interface | Contacts at Dynamic Interface |
|---|---|---|---|
| 8 | 1.47 | 0 | 0 |
| 38 | 3.40 | 3 | 0 |
| 39 | 3.03 | 8 | 1 |
| 41 | 1.91 | 23 | 3 |
| 44 | 2.67 | 3 | 6 |
| 130 | 2.52 | 2 | 0 |
| 259 | 2.62 | 0 | 9 |
| 264 | 3.34 | 0 | 32 |
| 470 | 3.76 | 0 | 0 |
| 496 | 3.04 | 0 | 18 |
| 543 | 3.94 | 0 | 0 |
| 565 | 3.98 | 0 | 0 |

**Supplementary Table 2. Plasmids used in this study.** For each plasmid, the table notes the name, resistance marker ($Ab^R$), origin type (ori), promoter (P), SiR-HP variant (SiR-HP), and domain inserted. With all domain insertion variants, identical linkers, GGGGSGGGGS, were used to fuse both the N- and C-termini of the ER-LBD to SiR-HP. All vectors express SiR-FP in addition to SiR-HP.

| Name | AbR | ori | P | SiR-HP | Domain inserted |
|---|---|---|---|---|---|
| pLW001 | CmR | ColE1 | $P_{tet}$ | wt SiR-HP | --- |
| pLW002 | CmR | ColE1 | $P_{tet}$ | wt SiR-HP | --- |
| pLW003 | CmR | ColE1 | $P_{tet}$ | SiR-HP R83A | --- |
| pDV8 | CmR | ColE1 | $P_{tet}$ | DV8 | ER-LBD |
| pDV38 | CmR | ColE1 | $P_{tet}$ | DV38 | ER-LBD |
| pDV39 | CmR | ColE1 | $P_{tet}$ | DV39 | ER-LBD |
| pDV41 | CmR | ColE1 | $P_{tet}$ | DV41 | ER-LBD |
| pDV44 | CmR | ColE1 | $P_{tet}$ | DV44 | ER-LBD |
| pDV130 | CmR | ColE1 | $P_{tet}$ | DV130 | ER-LBD |
| pDV259 | CmR | ColE1 | $P_{tet}$ | DV259 | ER-LBD |
| pDV264 | CmR | ColE1 | $P_{tet}$ | DV264 | ER-LBD |
| pDV470 | CmR | ColE1 | $P_{tet}$ | DV470 | ER-LBD |
| pDV496 | CmR | ColE1 | $P_{tet}$ | DV496 | ER-LBD |
| pDV543 | CmR | ColE1 | $P_{tet}$ | DV543 | ER-LBD |
| pDV565 | CmR | ColE1 | $P_{tet}$ | DV565 | ER-LBD |

### CITATIONS


1. Cepeda, M. R., McGarry, L., Pennington, J. M., Krzystek, J. & Stroupe, M. E. The role of extended Fe4S4 cluster ligands in mediating sulfite reductase hemoprotein activity. *Biochimica et Biophysica Acta (BBA) - Proteins and Proteomics* **1866**, 933–940 (2018).
2. Esfahani, B. G. *et al.* Structure of dimerized assimilatory NADPH-dependent sulfite reductase reveals the minimal interface for diflavin reductase binding. Preprint at https://doi.org/10.1101/2024.06.14.599029 (2024).
3. Askenasy, I. *et al.* The N-terminal Domain of Escherichia coli Assimilatory NADPH-Sulfite Reductase Hemoprotein Is an Oligomerization Domain That Mediates Holoenzyme Assembly. *Journal of Biological Chemistry* **290**, 19319–19333 (2015).
